# A Tethered Ligand Assay to Probe SARS-CoV-2:ACE2 Interactions

**DOI:** 10.1101/2021.08.08.455468

**Authors:** Magnus S. Bauer, Sophia Gruber, Adina Hausch, Lukas F. Milles, Thomas Nicolaus, Leonard C. Schendel, Pilar López Navajas, Erik Procko, Daniel Lietha, Rafael C. Bernardi, Hermann E. Gaub, Jan Lipfert

## Abstract

SARS-CoV-2 infections are initiated by attachment of the receptor-binding domain (RBD) on the viral Spike protein to angiotensin-converting enzyme-2 (ACE2) on human host cells. This critical first step occurs in dynamic environments, where external forces act on the binding partners and multivalent interactions play critical roles, creating an urgent need for assays that can quantitate SARS-CoV-2 interactions with ACE2 under mechanical load and in defined geometries. Here, we introduce a tethered ligand assay that comprises the RBD and the ACE2 ectodomain joined by a flexible peptide linker. Using magnetic tweezers and atomic force spectroscopy as highly complementary single-molecule force spectroscopy techniques, we investigate the RBD:ACE2 interaction over the whole physiologically relevant force range. We combine the experimental results with steered molecular dynamics simulations and observe and assign fully consistent unbinding and unfolding events across the three techniques, enabling us to establish ACE2 unfolding as a molecular fingerprint. Measuring at forces of 2-5 pN, we quantify the force dependence and kinetics of the RBD:ACE2 bond in equilibrium. We show that the SARS-CoV-2 RBD:ACE2 interaction has higher mechanical stability, larger binding free energy, and a lower dissociation rate in comparison to SARS-CoV-1, which helps to rationalize the different infection patterns of the two viruses. By studying how free ACE2 outcompetes tethered ACE2, we show that our assay is sensitive to prevention of bond formation by external binders. We expect our results to provide a novel way to investigate the roles of mutations and blocking agents for targeted pharmaceutical intervention.

## INTRODUCTION

A subset of coronaviruses (CoV) cause severe acute respiratory syndrome (SARS) in humans. We have seen three major outbreaks in recent years, including the first SARS pandemic from 2002-2004 (SARS-CoV-1), middle east respiratory syndrome (MERS-CoV) that first emerged in 2012, and the ongoing COVID-19 pandemic (SARS-CoV-2). SARS-CoV-2 particles carry on the order of 100 copies of the trimeric viral glycoprotein Spike (S) on their surface (1), giving the appearance of an eponymous corona around the virus. Like SARS-CoV-1, SARS-CoV-2 attaches to human host cells by S binding to angiotensin-converting enzyme-2 (ACE2) (2-6) (Fig. 1A). Specifically, each S trimer carries a receptor-binding domain (RBD), presented in an up or down conformation, at the tips of the three S1 subunits that can bind ACE2 in the up conformation (Fig. 1B) (7). Binding of the virus to host cells occurs in dynamic environments (8, 9) where external forces act on the virus particle. In particular, in the respiratory tract, coughing, sneezing, and mucus clearance exert mechanical forces (10, 11) that the virus must withstand for productive infection. However, the exact magnitude and dynamics of applied forces are unknown and likely variable.

**Fig. 1.**
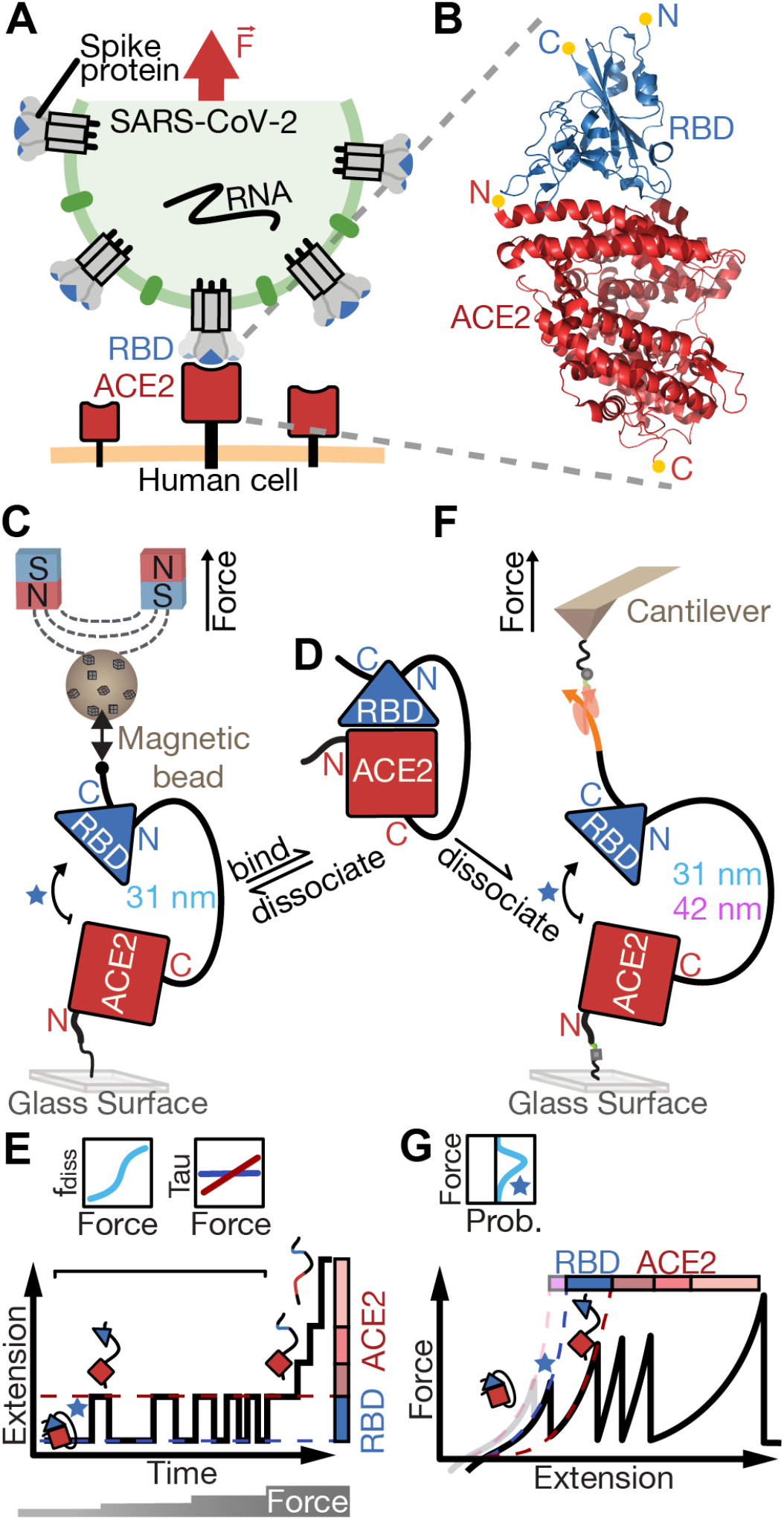
Single-molecule assays to probe the SARS-CoV-2 RBD:ACE2 interface under force. Motivation and overview of our tethered ligand assay for equilibrium and dynamic single-molecule force spectroscopy (SMFS) measurements in magnetic tweezers (MT) and atomic force microscopy (AFM). **A** Schematic of a SARS-CoV-2 virus particle (green) presenting spike protein trimers (grey) that can bind to human ACE2 (red) on the cell surface via their RBDs (blue). The bond between RBD and ACE2 is formed in a dynamic environment, where it must withstand external mechanical forces (indicated by the red arrow), e.g. caused by coughing or sneezing in the respiratory tract, in order to allow efficient infection of the human host cell (orange). **B** Crystal structure of the RBD:ACE2 complex (PDB ID: 6m0j). The N- and C-termini of the RBD (blue) and ACE2 (red) are indicated with yellow dots. **C** Schematic (not to scale) of the tethered ligand assay in MT. The tethered ligand construct consists of the ACE2 ectodomain (red square) and RBD (blue triangle) joined by a flexible polypeptide linker (black line) of 85 amino acids (31 nm contour length) or 115 amino acids (42 nm contour length). The tethered ligand construct is attached with one end covalently to the surface via an elastin-like polypeptide (ELP) linker (25) and with the other end to a superparamagnetic bead via a biotin-streptavidin bond. Permanent magnets above the flow cell enable the application of precisely calibrated stretching forces. **D** Tethered ligand construct in the absence of force, where the RBD remains bound to ACE2. **E** Stylized measurement of the tethered ligand construct in the MT. To probe RBD:ACE2 bond dynamics, time traces of the tether extension are recorded at different levels of applied force (indicated on the bottom). At low forces, reversible transitions between the bound configuration, with the RBD:ACE2 interface engaged, and an dissociated configuration, where the interface is broken and the peptide linker connecting the domains is stretched, are observed as jumps between two extension levels (red and blue dashed lines). At higher forces, further upward steps in the extension trace correspond to unfolding events of protein (sub-)domains. From the MT time traces, both the fraction of time spent in the dissociated state and the dwell times in the bound and dissociated state can be determined as a function of applied force (top). **F** Schematic (not to scale) of the tethered ligand construct in the AFM. Here, covalent attachment to the surface uses a heterobifunctional polyethylene glycol (PEG) spacer and the coupling to the AFM cantilever is accomplished via an Fgγ tag on the protein that binds with very high force stability to the ClfA protein handle on the cantilever. **G** Stylized AFM measurement. The cantilever is retracted with constant velocity and the force response to the applied extension is shown as a force-extension curve. With increasing extension the RBD:ACE2 interface ruptures (blue star), protein subdomains unfold, and finally the ClfA:Fgγ bond ruptures, giving rise to distinct peaks in the force-extension curve. Comparing two constructs with different linker lengths (31 nm black solid line and 42 nm grey/lilac alternative first peak) joining RBD and ACE2 allows assignment of the RBD:ACE2 interface rupture (blue star) and unfolding of parts of the RBD to the first increment. Histograms of rupture forces (top) are compiled from multiple measurements.

The SARS-CoV-2 S protein and its interaction with ACE2 have been the target of intense research activity, as they are critical in the first steps of SARS-CoV-2 infection and S constitutes a major drug and the key vaccine target in the current fight against COVID-19. Further, differences in binding between ACE2 and the SARS-CoV-1 and SARS-CoV-2 RBDs have been linked to the different observed patterns in lower and upper respiratory tract infections by the two viruses (5). Despite its importance, many questions about RBD:ACE2 interactions, particularly about their stability under external forces, are unresolved. Consequently, there is an urgent need for assays that can probe the affinity and kinetics of the interaction under a wide range of external forces. In nature, receptor-ligand pairs are often held in spatial proximity at high effective concentrations due to neighboring receptor: ligand interactions holding the viral membrane close to the cell surface. Conventional affinity measurements such as SPR and BLI do not take into account these effects. Therefore, there is a need for novel in vitro assays mimicking this when measuring bond characteristics. Here, we present a tethered ligand assay to determine RBD interactions with ACE2 at the single-molecule level subject to defined levels of applied force. Our assay utilizes fusion protein constructs of SARS-CoV-1 or SARS-CoV-2 RBD and the human ACE2 ectodomain joined by a flexible peptide linker. To probe the linkage under a large range of mechanical forces and loading rates, we used two highly complementary single-molecule force spectroscopy (SMFS) approaches: an atomic force microscope (AFM) and magnetic tweezers (MT) (Fig. 1C - G). We complemented the experiments with steered molecular dynamics (SMD) simulations to provide microscopic insights that are inaccessible experimentally.

AFM force spectroscopy can probe molecular interactions and protein stability dynamically (12-14), typically measuring at constant loading rate, and can investigate even the most stable high force host-pathogen interactions (at forces *F* > 1,000 pN) (15). In AFM experiments, the molecular construct of interest is stretched between a surface and the tip of an AFM cantilever. The cantilever is retracted at a constant velocity, and the force is monitored from the cantilever deflection. Molecular rupture or protein (sub-)domain unfolding events give rise to a sawtooth-like pattern in the force vs. extension traces (Fig. 1G). In contrast, MT typically operate at constant force and can resolve very low forces (16, 17), down to F < 0.01 pN. In MT, molecules are tethered between a flow cell surface and magnetic beads. External magnets apply defined and constant stretching forces and the tether extension is monitored by video microscopy. In the MT, unbinding or unfolding events give rise to steps in the extension vs. time trace (Fig. 1E).

Tethered ligand assays have provided insights into a range of critical molecular interactions under mechanical load (18-21). Under constant force, they allow observation of repeated interactions of the same binding partners, which are held in spatial proximity under mechanical control. Therefore, they can provide information on affinity, avidity, on- and off-rates, and mechanical stability (18, 20). Conversely, AFM force spectroscopy can perform dynamic measurements in a highly automated fashion and can reveal characteristic protein unfolding patterns, which can serve as molecular fingerprints (22) to select only properly folded and assembled molecular constructs for further analysis.

Probing our tethered ligand construct by AFM force spectroscopy, we reveal the dynamic force stability of the assembly. In combination with SMD simulations, we assign the increments revealed by force spectroscopy and establish the ACE2 unfolding pattern as a molecular fingerprint to select properly assembled tethers. Using MT, we measure the on- and off-rates at different levels of mechanical load and extrapolate to the thermodynamic stability at zero load. We compare the stability of the SARS-CoV-1 and SARS-CoV-2 RBD:ACE2 interactions in all three assays and consistently find a lower force stability for SARS-CoV-1 across the different techniques.

## RESULTS

### A tethered ligand assay to probe viral attachment under physiological forces using MT and AFM

We designed tethered ligand fusion proteins that consist of the SARS-CoV-1 RBD or SARS-CoV-2 RBD and the ectodomain of human ACE2 joined by flexible polypeptide linkers (Fig. 1E). Protein constructs were designed based on the available crystal structures (23, 24) of the SARS-COV-1 or SARS-COV-2 RBDs in complex with human ACE2 and carry short peptide tags at their termini for attachment in the MT and the AFM (Materials and Methods). For MT experiments, the protein constructs were coupled covalently to the flow cell surface via elastin-like polypeptide (ELP) linkers (25) and to magnetic beads via a biotin:streptavidin linkage (26). Tethering multiple proteins in MT enables parallel measurements of multiple molecules over extended periods (hours to weeks) at precisely controlled forces (26). In MT, bead positions and, therefore, tether extensions are tracked by video microscopy with ∼1 nm spatial resolution and up to kHz frame rates (27-29). For AFM experiments, we employ the same tethered ligand construct as used in the MT assay and covalently anchor it to a glass surface using PEG spacers. The key difference to the MT measurement is the use of an Fgγ tag on the protein together with ClfA as a reversible handle system instead of the biotin:streptavidin linkage to attach it to the AFM cantilever. The ClfA:Fgγ interaction is non-covalent, but can withstand extremely high forces of up to 2 nN, making it a reliable attachment modality with a built-in force fingerprint for AFM force spectroscopy (15). Together with stable custom-built AFM setups (30), this enables reliable recordings of specific force-extension traces over several days.

### Dynamic AFM force spectroscopy reveals a characteristic unfolding pattern

The AFM traces of the tethered ligand constructs feature a total of five sawtooth-like peaks, each corresponding to an unfolding or unbinding event (Fig. 1G, Fig. 2A, and Supplementary Fig. S1). The last (right-most) peak exhibits forces well within the range previously established for the ultra-strong ClfA:Fgγ interaction (15), clearly indicating specific attachment of the protein construct. Consequently, the four peaks at lower forces must correspond to unfolding and unbinding events in a single tethered ligand construct. To visualize the most probable force-extension trace, we aligned and superimposed all individual curves (22, 31) (Fig. 2B and C). They all feature the final rupture peak assigned to the ClfA:Fgγ linkage and an initial (left-most) peak typically at lower forces around 26 pN for the SARS-CoV-1 fusion construct and about twice as high forces for the SARS-CoV-2 fusion construct (around 57 pN; Fig. 2 A on the right). This initial peak is followed by a trident-shaped, three-peak pattern at around 40 pN. The released contour lengths corresponding to each of these unbinding/unfolding peaks were determined from contour length transformations (22, 32) of each specific curve (see Materials and Methods). The four increments show the same order and very similar lengths for the SARS-CoV-1 and SARS-CoV-2 constructs (Fig. 2B,C, top insets).

**Fig. 2.**
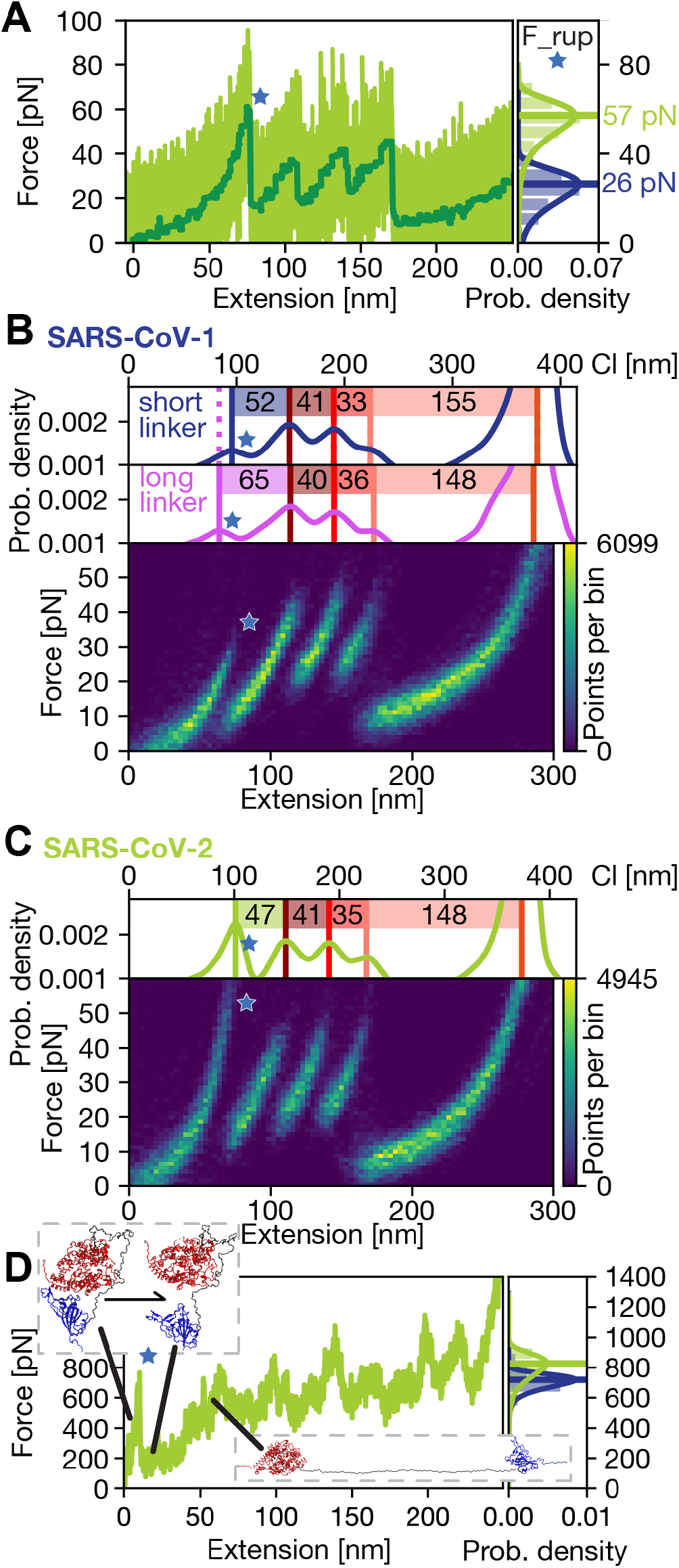
AFM force spectroscopy reveals multiple defined transitions in the tethered ligand construct. **A** AFM force-extension trace of the SARS-CoV-2 RBD:ACE2 tethered ligand construct (raw data in light green, total variation denoised dark green, recorded at 800 nm/s retraction velocity). Four defined peaks are clearly visible between 30 pN and 60 pN. The final ClfA:Fgγ rupture at > 1500 pN is not shown for clarity. **B** Heatmap of unfolding curves for the SARS-CoV-1 RBD:ACE2 fusion construct generated from 151 unfolding traces similar to the measurement shown in A. **C** Heatmap of unfolding curves for the SARS-CoV-2 RBD:ACE2 fusion construct generated from 127 unfolding curves, including the curve shown in **A**. Overall, the unfolding patterns look very similar for SARS-CoV-1 and 2, the only clear difference is that the first peak (indicated by a blue star throughout) is at higher forces for SARS-CoV-2. The right inset in panel **A** directly compares the most probable rupture forces determined by the Bell-Evans model for the first peak for the two constructs. The insets in the top of panels **B** and **C** show transformations of the force-extension data to contour length space (Materials and Methods). For SARS-CoV-1 two different length linkers were measured (31 nm linker in blue and 42 nm in purple); the difference of 13 nm in contour length for the initial peak is very close to the expected value. **D** Force extension trace obtained by a steered molecular dynamic (SMD) simulation of the SARS-CoV-2 RBD:ACE2 fusion protein shows qualitatively similar unfolding behavior as the AFM curve. Small renderings illustrate the course of the unfolding. The right inset in panel **D** compares the rupture forces of the RBD:ACE2 interface of SARS-CoV-1 and SARS-CoV-2 from multiple SMD simulations. The rupture forces in the SMD simulations are much higher than for the AFM measurements, due to the much higher loading rates (2.5 Å/ns vs. 800 nm/s). However, similar to AFM measurements, the simulations suggest that SARS-CoV-1 binding has lower force stability compared to SARS-CoV-2.

The fact that the first peak (Fig. 2, blue star) has a notably different force signature for the SARS-CoV-1 and SARS-CoV-2 constructs, while all other peaks are very similar, suggests that the first peak involves the RBD domain, which is the only part that is different between the two constructs. To probe whether the first peak is due to the RBD:ACE2 interface opening or RBD unfolding, we performed control experiments with a longer linker (115 amino acids (aa) corresponding to 42 nm contour length, instead of 85 aa or 31 nm) between the two domains. The measurements with the longer linker reveal an increase in the contour length released in the first peak by 13 nm. This is very close to the expected 11 nm and strongly suggests that the first peak represents dissociation of the RBD:ACE2 interface (Fig. 2B). The contour length increments of the first peaks (52 nm or 65 nm for the short and long linker, respectively), are, however, too large to be only caused by the interface opening, releasing the linker lengths (31 nm or 42 nm) and inducing a reorientation of the domains. Conversely, full unfolding of the RBD domain would release 193 aa or 70 nm contour length, much longer than the increments that are experimentally observed. Therefore, the first peak must involve interface opening and partial unfolding of the RBD. Since all measurements were conducted under non-reducing conditions (TBS buffer, see Materials and Methods), we expect disulfide bridges between cysteines to be formed, which shield large parts of the RBD structure from force (24, 33) and allow only 51 aa to unfold, corresponding to 19 nm contour length, in excellent agreement with experimentally observed increments (Supplementary Fig. S2).

With the first peak assigned to the RBD:ACE2 interface opening and partial RBD infolding, the subsequent trident-shaped, three-peak pattern is likely due to (step-wise) unfolding of the ACE2 domain. Control measurements with the ACE2 domain only and the same affinity tags resulted in traces showing the same trident-shaped pattern and no additional first peak (Supplementary Fig. S3), confirming the assignment of the trident-shaped pattern to the ACE2 domain. In conclusion, the first rupture event in the AFM measurements corresponds the RBD:ACE2 interface combined with partial RBD unfolding. Comparing the same constructs with the SARS-CoV-1 and the SARS-CoV-2 RBD tethered to ACE2, reveals a lower force stability for the SARS-CoV-1 interface compared to SARS-CoV-2 (Fig. 2A, histograms). In addition, the AFM data suggest that, after the opening of the interface, step-wise ACE2 unfolding gives rise to a defined pattern that can be used as a molecular fingerprint.

### All-atom steered molecular dynamic (SMD) simulations provide insights into the unfolding patterns in molecular detail

In an *in silico* SMFS approach, the tethered ligand protein probed in the AFM measurement was modeled using QwikMD (34). Based on the available crystal structures, RBD:ACE2 was modeled with and without the polypeptide linkers for both SARS-CoV-1 and SARS-CoV-2, in a total of 4 different systems (Materials and Methods). Over 300 SMD simulations were performed employing GPU-accelerated NAMD 3 (35). In the simulations, the behavior of the complexes agrees with the AFM experiments, revealing an unfolding starting with the dissociation of the RBD domain from ACE2 and consecutive unfolding of the RBD (Fig. 2D). The unfolding of parts of the RBD is caused by the linker that gets stretched after the interface is released from ACE2. After the initial RBD unfolding, ACE2 unfolds in several substeps. The corresponding force-distance curve agrees with the observed unfolding pattern obtained by AFM measurements. The insets in Fig. 2D show renderings at two different time points to visualize the forced RBD:ACE2 dissociation observed in SMD and AFM experiments.

Simulations of the RBD and ACE2 without linker showed identical behavior until the point of the interface rupture and the rupture process was conserved for the protein models with and without the linker. In this way the rupture forces of SARS-CoV-1 and SARS-CoV-2 could be determined both for the tethered ligand protein and for just the single domains. For collecting statistics, the construct without linker was used to determine unbinding forces of the RBD:ACE2 interface. The simulations give 20% higher forces for SARS-CoV-2 compared to SARS-CoV-1 (Fig. 2D, right inset). The observed higher forces for SARS-CoV-2 qualitatively agree with the differences in forces determined from the AFM measurements with an expected increase in absolute forces due to the much higher loading rates in the SMD simulation compared to SMFS.

### Unfolding patterns across different force-loading rate regimes are highly reproducible

In contrast to AFM and SMD simulations, where tethered fusion constructs were subjected to forces with constant force-loading rate, MT were used to examine the fusion complex at constant forces, first studying equilibrium binding and dissociation between RBD and ACE2 at lower forces and then unfolding individual protein domains at high forces (Fig. 3A). At forces between 2 to 5 pN, we observe systematic transitions in the extension traces, with jumps between a high extension “dissociated” and low extension “bound” state (Fig. 3A, Equilibrium and Fig. 4A,B). The transitions systematically shift towards the dissociated state with increasing force (Fig. 4A). Alternating between levels of low force (0.5 pN) and higher forces (15, 20, 25, and 30 pN) (Fig. 3A, “High Forces”) reveals three distinct unfolding transitions. At forces of 15 and 20 pN, one unfolding transition repeatedly occurs after refolding during a low force interval. This reversible transition corresponds to the systematic transitions recorded at equilibrium between 2 and 5 pN (Fig. 3A blue boxes and Fig. 3B, blue histogram for high force transitions), while a subsequent two-step transition above 25 pN (Fig. 3A, red box and Fig. 3B, red histograms) is irreversible. Measuring just the ectodomain of ACE2 revealed the same characteristic irreversible two-step unfolding pattern above 25 pN without showing the systematic transitions at lower forces or the reversible unfolding at 15 and 20 pN (Supplementary Fig. S5). This suggests a clear assignment of the two irreversible unfolding events to the unfolding of ACE2, while the equilibrium transitions, observed as reversible steps at 15 and 20 pN are attributed to the RBD:ACE2 interface opening and partial RBD unfolding, corresponding to the first peak in the AFM force-distance curves. Using the worm-like-chain model (36) with a persistence length of 0.5 nm (26) the contour length of the observed extension increments in MT are calculated and a Gaussian distribution is fitted to the histograms (Fig. 3B). The mean values of the increments in MT (Fig. 3C, MT) are compared to the contour length transformed increments in the AFM (Fig. 3C, AFM). For an overview of the contour length transformed increments in MT and AFM refer to Table 1. Strikingly, the increments observed with both single-molecule force spectroscopy techniques are in excellent agreement, within experimental errors, with each other and with theoretical expectations based on the crystal structure (24) and the fusion construct design. Comparing the total length gain from unfolding the tethered ligand ACE2 in the AFM (224 nm) and MT (233 ± 25.5 nm), reveals quantitative agreement within experimental errors. The first ACE2 unfolding event (Table 1, ACE2_1) in the AFM almost perfectly matches the first increment in MT. The second and third ACE2 unfolding (Table 1 ACE2_2 and ACE2_3) in the AFM cannot be separately resolved in MT for most molecules and typically occur as one single large step (Table 1 ACE2_combined). In a small subpopulation (8 of 42 total molecules), however, we observed a very short-lived intermediate level, splitting the large step into a smaller and yet another larger step, matching the increments observed in the AFM. In summary, ACE2 unfolding provides a highly specific and reproducible molecular fingerprint across AFM and MT measurements that we subsequently used to select for specific tethers to probe the RBD:ACE2 interface.

**Table 1.** Increments of high-force transitions in MT and unfolding peaks in the AFM of the SARS-CoV-2 RBD:ACE2 tethered ligand construct. Data are mean and std for 42 molecules in MT and 127 molecules in the AFM. In MT, the ΔACE2_2 and ΔACE2_3 are observed separately only in a small sub-population (8 out of 42 molecules). Mostly they are combined into one step ΔACE2_combined. The large error for the smaller intermediate step is due to imprecisions in the increment measurement due to the short lifetime of this state.

|  | MT increments (nm) | AFM increments (nm) |
| --- | --- | --- |
| $\Delta\text{RBD}$ | $50.7 \pm 10.8$ | 47 |
| $\Delta\text{ACE2\_1}$ | $42.1 \pm 5.5$ | 41 |
| $\Delta\text{ACE2\_2}$ | $45.7 \pm 14.0$ | 35 |
| $\Delta\text{ACE2\_3}$ | $147.6 \pm 7.8$ | 148 |
| $\Delta\text{ACE2\_combined}$ | $191.3 \pm 18.7$ | -- |

**Fig. 3.**
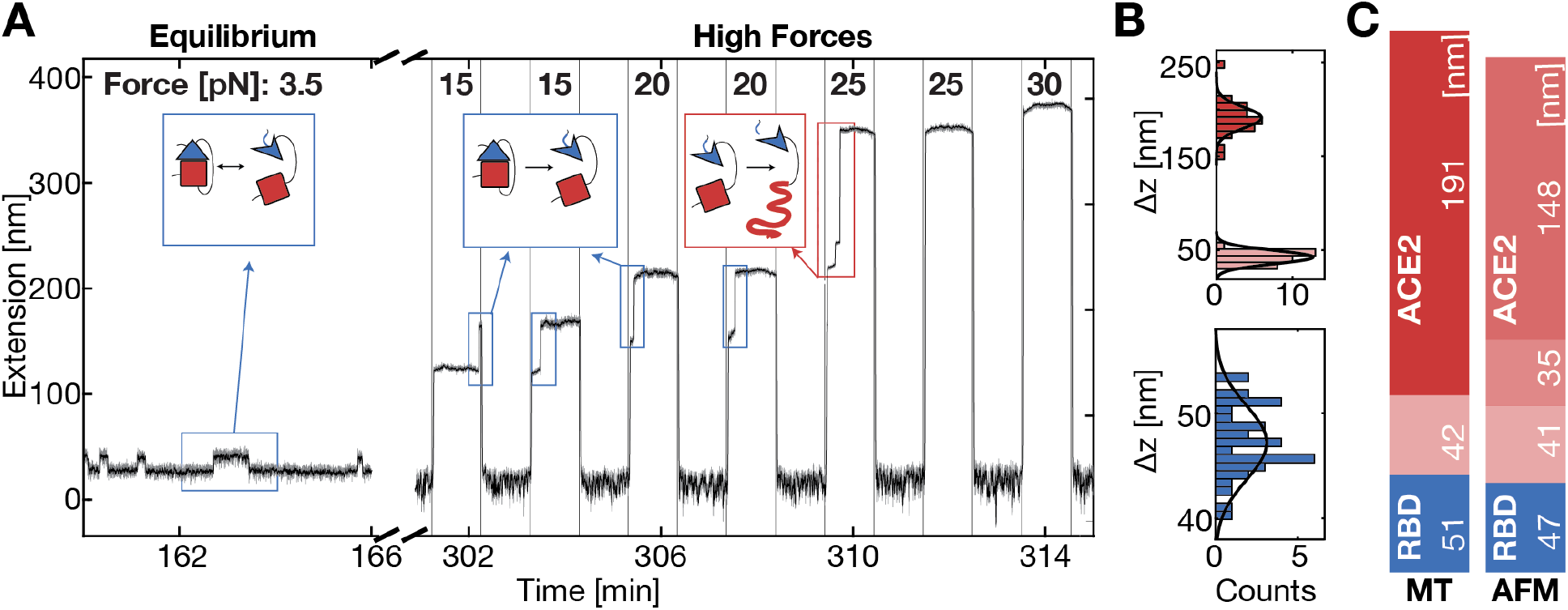
SARS-CoV-2 RBD:ACE2 interface opening and unfolding in MT. **A** Extension time trace of an RBD:ACE2 tethered ligand construct in MT shows distinct transitions at different levels of constant force. At low forces (labelled “Equilibrium”), we observe stochastic transitions between two extension levels, separated by ΔRBD ≈ 13.5 nm at a force of 3.5 pN. In subsequent measurements at higher forces (labelled “High Forces”), the force is iteratively altered between 0.5 pN (low extension) and increasingly higher forces (indicated at the top). At 15 and 20 pN, the RBD:ACE2 interface ruptures together with a partial RBD unfolding (ΔRBD ≈ 32 nm) and refolds in the subsequent 0.5 pN interval. At 25 pN, ACE2 irreversibly unfolds in two steps, first a smaller one (ΔACE2small ≈ 30 nm) and then a larger one (ΔACE2large ≈ 135 nm). Grey trace: 5-frame moving average filtered; Black trace: 40-frame moving average filtered. **B** Histograms of unfolding increments at high forces of the RBD (blue) and the two parts of ACE2 (pink, red) in MT. The histograms are fitted with a Gaussian (solid black line). Some points are outside of the plotting range for clarity but included in the fit. **C** Mean unfolding increments observed for tethered ligand construct with constant forces in MT and with constant loading rate in AFM. The observed increments were transformed into contour lengths using a WLC model, assuming a persistence length of 0.5 nm in MT and 0.365 nm in the AFM. Values ± standard deviation for the increments in MT are: ΔRBD = (51 ± 10.8) nm, ΔACE2_1 = (42 ± 5.5) nm, Δ ACE2_combined = (191 ± 18.7) nm. Values for the increments in the AFM are: ΔRBD = 47 nm, ΔACE2_1 = 41 nm, Δ ACE2_2 = 35 nm, and ΔACE2_3 = 148 nm. Values in **B** and **C** correspond to mean values from 42 molecules observed in MT and 127 molecules observed in AFM.

**Fig. 4.**
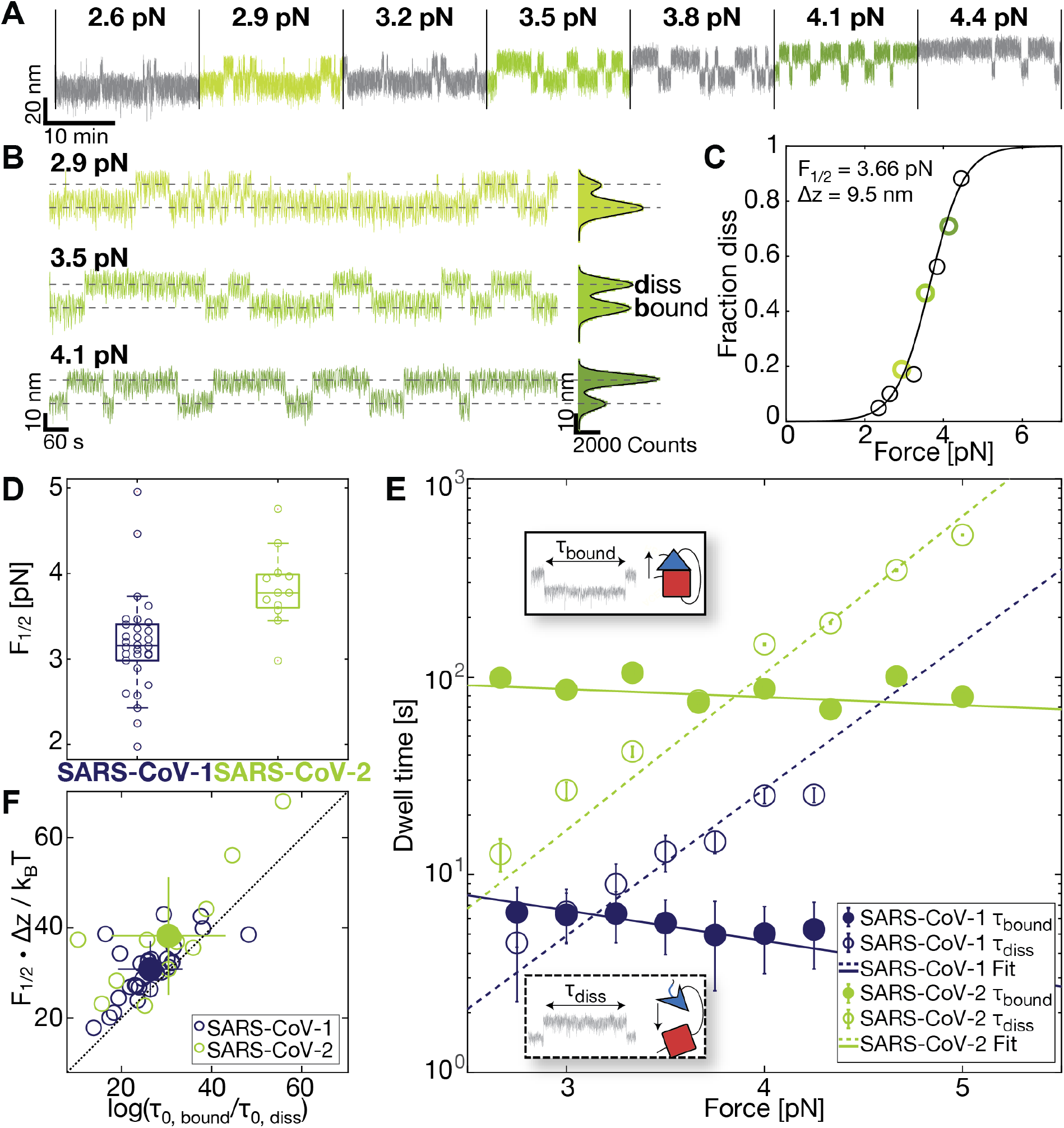
Comparison of mechanical stability and kinetics of ACE2 binding to SARS-CoV-1 and SARS-CoV-2 RBD. **A** Extension time traces at different constant forces for the SARS-CoV-2 RBD:ACE2 fusion construct reveal stochastic transitions between two extension levels, corresponding to the bound and dissociated RBD:ACE2 interface, respectively. Increasing the force increases the fraction of time spent in the dissociated conformation. **B** Expanded views of extensions at three different forces: below, at, and above the mid-force from trace in **A** show shift towards the dissociated conformation with increasing force. Same color code as panel **A**. The “dissociated” and “bound” states are indicated by dashed lines. The histograms of the extensions are fitted with a double Gaussian (solid black lines). **C** The fraction of time in the dissociated conformation determined from extension time traces (symbols; points determined from the traces in panel **B** are shown with matching color codes). The black line is a fit of Equation 1. Fitting parameters *F*_1/2_ and Δx are shown as an inset. **D** Comparison of the *F*_1/2_ distribution between SARS-CoV-1 and SARS-CoV-2 reveal a significantly higher force stability of the SARS-CoV-2 RBD:ACE2 bond (p = 0.00124; two-tailed p-test). Data point are the fitted *F*_1/2_ from independent molecules. Boxes are the median and interquartile range. **E** Dwell times in the dissociated (open symbols) and bound state (filled symbols) determined from extension time traces for SARS-CoV-1 (blue) and SARS-CoV-2 (green). Mean dwell times for individual molecules were determined from maximum likelihood fits of a single exponential to the dwell time distributions. The symbols and error bars are the mean and standard deviation from log-averaging over 29 (SARS-CoV-1) and 12 (SARS-CoV-2) molecules. Dashed and solid lines correspond to the mean of the exponential fits to the individual dwell times in the bound state and dissociated state, respectively. The insets visualize dwell times in MT time-extension traces. **F** Free energy differences between the bound and dissociated state of the RBD:ACE2 tethered ligand constructs. The free energy differences were obtained from the equilibrium data as *F*_1/2_·Δx and from the dynamics as log(τ_0,bound_ / τ_0,diss_). The data fall along the 45 degree line (dashed), indicating that the two estimates give consistent values. Comparison of the SARS-CoV-1 and SARS-CoV-2 data reveals a larger free energy difference for SARS-CoV-2, indicating a more stable interface. Distributions and mean values shown in panel **D - F** are for 29 (SARS-CoV-1) and 12 (SARS-CoV-2) molecules, respectively.

### MT measurements probe the RBD:ACE2 interaction under load in equilibrium

After selecting specific tethers based on the ACE2 fingerprint, we analyze the equilibrium transitions measured in MT at forces between 2 and 5 pN. The transitions systematically change with applied force: At low forces, the interface is predominantly formed (bound state), while increasing the force increases the fraction of time with an open interface and a partially unfolded RBD (dissociated state) (Fig. 4A). Histograms of the tether extension revealed two clearly separated peaks that are fit well by a double Gaussian (Fig. 4B; bold black lines). By setting thresholds at the minimum between the extension peaks, we defined populations in the dissociated and bound states. The fraction in the dissociated state (Fig. 4C; circles), follows a sigmoidal force dependence. The data are well-described by a two-state model (Fig. 4C; solid line) where the free energy difference between the two states (bound vs. dissociated but still attached to each other with a linker) depends linearly on the applied force F, i.e. ΔG = ΔG_0_ –F·Δz, such that the fraction in the dissociated state is

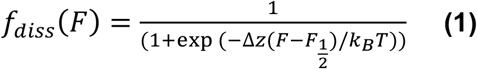

where k_B_ is Boltzmann’s constant and T the absolute temperature. *F*_1/2_ and Δz are fitting parameters that represent the midpoint force, where the system spends half of the time in the dissociated and half of the time in the bound conformation, and the distance between the two states along the pulling direction, respectively. For the two states of the fusion protein, the free energy difference at zero force is given by ΔG_0_ = *F*_1/2_·Δz and provides a direct measure of the stability of the binding interface.

From fits of Equation 1 to the data for the SARS-CoV-2 RBD:ACE2 construct, we found F_1/2_ = 3.8 ± 0.4 pN and Δz = 10.2 ± 3.7 nm, and, therefore, ΔG_0_ = F_1/2_·Δz = 5.5 ± 2.1 kcal/mol (data are the mean and standard deviation from fits to biological repeats; see Table 2 for a summary of all fitted parameters). The value of Δz determined from fitting Equation 1 is within experimental error in agreement with the distance between the dissociated and bound states Δz_G_= 13.5 ± 1.8 nm determined from fitting two Gaussians to the extension histograms at the equilibrium force *F*_1/2_ and evaluating the distance between the fitted center positions. The observed Δz is also in excellent agreement with the expected extension change of ≈ 13.4 nm, based on the crystal structure (24) (PDB ID: 6m0j) taking into account the stretching elasticity of the 85 aa protein linker and the unfolding of non-shielded parts of the RBD using the worm-like chain (WLC) model (26, 37, 38) with a bending persistence length of L_p_ = 0.5 nm and assuming 0.365 nm/aa.

**Table 2.** Interaction parameters for ACE2 and SARS-CoV-2 or SARS-CoV-1 RBD determined using the tethered ligand assay. Data are the mean and std from N = 12 and 29 molecules, respectively.

|  | SARS-CoV-2:<br>ACE2-linker-RBD<br>(85 aa linker) | SARS-CoV-1:<br>ACE2-linker-RBD<br>(85 aa linker) |
| --- | --- | --- |
| $F_{1/2}$ | $3.8 \pm 0.4$ pN | $3.2 \pm 0.6$ pN |
| $\Delta z$ (from fit of Equation 1) | $10.2 \pm 3.7$ nm | $9.7 \pm 1.7$ nm |
| $\Delta z_G$ (from fit of two Gaussians) | $13.5 \pm 1.8$ nm | $11.3 \pm 1.7$ nm |
| $\Delta G_0 (= \Delta z \cdot F_{1/2})$ | $5.5 \pm 2.1$ kcal/mol | $4.4 \pm 1.1$ kcal/mol |
| $\tau_{0,\text{diss}}$ | $0.07 \pm 0.19$ s | $0.03 \pm 0.05$ s |
| $\tau_{0,\text{bound}}$ | $114.5 \pm 278.8$ s | $19.0 \pm 24.8$ s |
| $\Delta G_{0,\text{tau}} (= k_B T \cdot \log(\tau_{0,\text{diss}}/\tau_{0,\text{bound}}))$ | $4.4 \pm 1.7$ kcal/mol | $3.8 \pm 2.0$ kcal/mol |
| $\Delta z_{\text{diss}}$ | $7.5 \pm 2.6$ nm | $7.0 \pm 1.8$ nm |
| $\Delta z_{\text{bound}}$ | $0.4 \pm 2.5$ nm | $1.5 \pm 1.6$ nm |

In addition to providing information on the binding equilibrium, the MT assay gives access to the binding kinetics under force. Analyzing the extension time traces using the same threshold that was used to determine the fraction dissociated vs. force, we identify dwell times in the dissociated and bound states (Supplementary Fig. S4A, B), which are exponentially distributed (Supplementary Fig. S4C, D). The mean dwell times in the dissociated state increase with increasing force, corresponding to the intuitive interpretation that the higher the force, the longer it takes for a dissociated receptor:ligand pair to rebind (Fig. 4E, dashed green line). The mean dwell times in the bound state, on the other hand, decrease with increasing force, albeit only slightly, implying that the higher the force, the shorter the bond stays intact. The dependencies of the mean dwell times on the applied force F are well described by exponential, Arrhenius-like relationships (39)

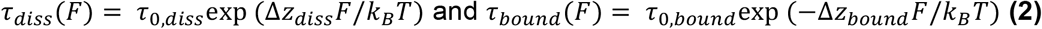

where the fitting parameters τ_0,diss_ and τ_0,bound_ are the lifetimes of the dissociated and bound conformation in the absence of force and Δz_diss_ and Δz_bound_ are the distances to the transition state along the pulling direction.

The parameters Δz_diss_ and Δz_bound_ quantify the force-dependencies of the lifetimes of the respective states, and the slopes in the log(τ_diss/bound_) vs. F plots (Fig. 4E) are given by Δz_diss/bound_ / k_B_T. Δz_bound_ is smaller than Δz_diss_ (by more than a factor of ∼ 5), i.e. dissociation of the bound complex is less force-sensitive than rebinding from the dissociated conformation. The different force sensitivities can be rationalized from the underlying molecular processes: The bound complexes feature protein-protein interactions that will break over relatively short distances; in contrast, the dissociated conformations involve flexible peptide linkers that make rebinding from the dissociated states more force-dependent.

The extrapolated lifetimes at zero force of the bound conformations τ_0,bound_ are in the range of 115 s for SARS-CoV-2. In comparison, the lifetimes of the dissociated states in the absence of load τ_0,diss_ are much shorter, in the range of ∼ 0.07 s (Table 2). The extrapolated lifetimes at zero force provide an alternative route to computing the free energy difference between the previously described dissociated and bound states at *F* = 0, which is given by ΔG_0,tau_ = k_B_T·log(τ_0,diss_/τ_0,bound_). We find good agreement, within experimental error, between the free energy differences ΔG_0,tau_ determined from the extrapolated lifetimes and the values ΔG_0_ = F_1/2_·Δz from Equation 1 (Table 2, Fig. 4F), providing a consistency check between equilibrium and kinetic analysis. The results show that studying our tethered ligand assay in MT can yield consistent information on the binding equilibrium and on the interaction kinetics under external force.

### SARS-CoV-2 attachment is more stable and longer-lived than SARS-CoV-1 under constant load

A construct using the same 85 aa linker and attachment geometry, but the SARS-CoV-1 RBD instead of SARS-CoV-2 RBD, shows a qualitatively very similar force-response in MT, with stochastic transitions between a bound and a dissociated conformation at an equilibrium force below 5 pN and ACE2 unfolding at forces higher than 25 pN. The increments from unfolding the tethered ACE2 are in excellent agreement both with the increments from the ACE2 unfolding of the SARS-CoV-2 tethered ligand construct in MT and from the AFM ACE2 unfolding (Supplementary Fig. S5). Furthermore, they happened at comparable forces as in the SARS-CoV-2 tethered ligand construct and single ACE2 in MT. As for the SARS-CoV-2 tethered ligand construct, the molecules for equilibrium and kinetic analysis of the SARS-CoV-1 RBD:ACE2 bond could thus be selected based on the molecular fingerprint of ACE2 unfolding and directly compared to SARS-CoV-2 tethered ligand constructs.

From fits of Equation 1, we found F_1/2_ = 3.2 ± 0.6 pN, Δz = 9.7 ± 1.7 nm and thus ΔG_0_ = 4.4 ± 1.1 kcal/mol for the SARS-COV-1 RBD:ACE2 tethered ligand construct (Table 2). The length increment Δz is again, within experimental error, in good agreement with the value determined from fitting two Gaussians to the extension histogram near the midpoint of the transition (Δz_G_ = 11.3 ± 1.7 nm at *F*_1/2_). The slightly shorter extension increment upon dissociation for the SARS-CoV-1 construct compared to SARS-CoV-2, despite using the same 85 aa linker and a very similar crystallographic geometry is mostly due to the smaller extension of the WLC at the lower midpoint force for SARS-CoV-1. Comparing the two lengths after contour length transformation yields Δz_SARS-CoV-1 RBD, WLC_ = 51.2 ± 7.9 nm and Δz_SARS-CoV-2 RBD, WLC_ = 50.3 ± 7.7 nm, in agreement, within error. In contrast, the SARS-CoV-1 RBD:ACE2 interface shows a significantly lower midpoint force *F*_1/2_ than the SARS-CoV-2 RBD:ACE2 bond (Fig. 4D; *p* = 0.0029 from a two-sample *t*-test), in line with the lower unbinding forces for SARS-CoV-1 observed in the AFM (Fig. 2B, C) and SMD simulations. The difference also manifests in a reduced mean free energy of the SARS-CoV-1 RBD:ACE2 bond computed both from the midpoint forces and distances as well as from the rates (Fig. 4F). Comparing the lifetimes of the bound and dissociated states (Fig. 4E, blue and green lines) and extrapolating the mean lifetimes to zero force yields no significant difference between the lifetimes in the dissociated conformation (Fig. 4E, dashed lines, τ_0,diss, SARS-CoV-1_ = 0.03 ± 0.05 s, τ_0,diss, SARS-CoV-2_ = 0.07 ± 0.19 s, p = 0.293). In contrast, the extrapolated lifetime in the bound state at zero force, on the other hand, is significantly different at α = 0.1 (τ_bound, SARS-CoV-1_ = 19.0 ± 24.8 s, τ_bound, SARS-CoV-2_ = 114.5 ± 278.8 s, p = 0.072) and more than six times higher for SARS-CoV-2 RBD, indicating a lower dissociation rate from ACE2. Furthermore, the lifetime of the SARS-CoV-2 RBD:ACE2 bond decreases less with force, i.e. has a shallower slope than the bond lifetime of SARS-CoV-1 RBD:ACE2. Therefore, the longer lifetime of the SARS-CoV-2 bond compared to SARS-CoV-1 becomes even more pronounced under mechanical load.

### Magnetic tweezers provide a sensitive assay to study molecules that block the RBD:ACE2 interaction

Apart from providing a tool to assess equilibrium binding and kinetics, investigation of the tethered ligand assay in MT also allows us to probe the influence of other binding partners on the bond dynamics. For this purpose, the equilibrium bond dynamics are first recorded under standard conditions before exchanging the buffer and conducting the same experiment in the presence of the compound of interest. As a proof-of-concept, we investigate the influence of soluble ACE2 on the bond dynamics (Fig. 5). First, we recorded the equilibrium binding of the SARS-COV-2 RBD and the tethered ACE2 in a short measurement between 3.0 and 4.5 pN, observing the characteristic transitions between the dissociated and bound conformation (Fig. 5A, “Without free ACE2”). We then added 3.8 µM free, soluble ACE2 and increased the force to 7 pN, to ensure the dissociation between the tethered receptor-ligand pair and enable binding of free ACE2, before conducting the same measurement in the presence of the free ACE2 (Fig. 5A, “With free ACE2”). At the same forces, the system is now predominantly in the dissociated conformation, rebinding only occasionally. This matches the interpretation of free ACE2 binding to the RBD and thus preventing the tethered ACE2 from binding and transitioning into the bound conformation. Overall, we find that the number of dissociation and rebinding events is significantly reduced in the presence of free ACE2 (Fig. 5B, *p =* 0.022 from a repeated measures t-test over 6 independent molecules).

**Fig. 5.**
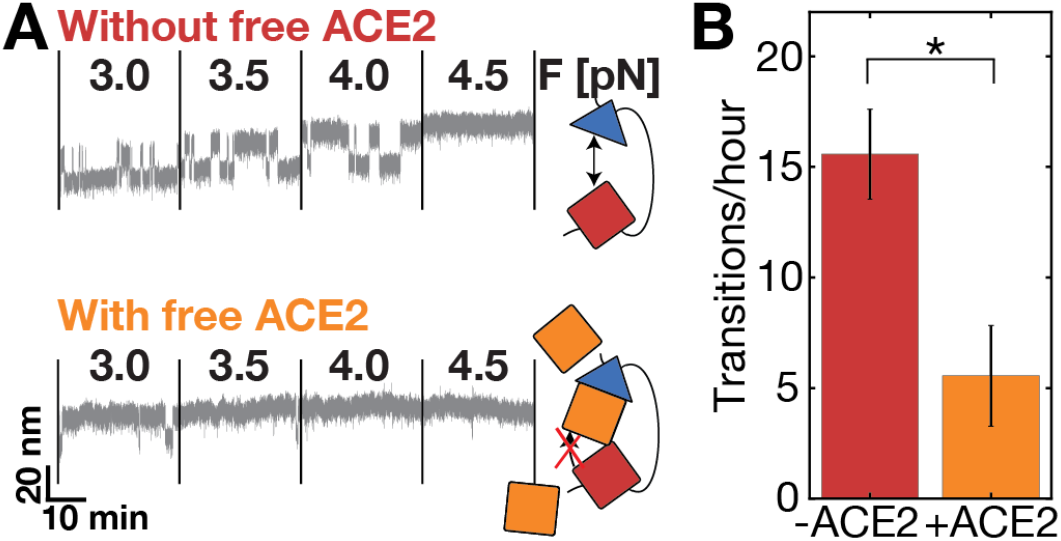
Blocking of ACE2:SARS-CoV-RBD bond with free ACE2. **A** Extension-time traces of a tethered ligand construct in MT, first without and then in the presence of free ACE2 (∼ 3.8 µM) in solution. Without free ACE2, the RBD:ACE2 bond frequently dissociates and rebinds (corresponding to changes in extension level). In the presence of free ACE2, the bond is significantly longer dissociated, rebinding only a few times (corresponding to a longer time spent at the level of high extension). **B** Quantification of mean number of transitions per hour with and without free ACE2 for 6 independent molecules. Error bars indicate standard error of the mean. The difference is statistically significant with p = 0.0218 (two-tailed t-test for two dependent means).

## DISCUSSION

In order to infect human host cells, SARS coronaviruses engage with receptors in the turbulent and dynamic environment of the respiratory tract, where forces impact their anchoring. Force-stability of the RBD:ACE2 bond thus plays a crucial role for the virus to be able to infect host cells efficiently. Understanding the binding mechanism and being able to assess the stability of the RBD:ACE2 bond opens opportunities to develop and test drugs and predict fitness advantages of mutated RBDs. Here, we report a tethered ligand assay to study and characterize the stability and binding mechanism of the SARS-CoV RBD:ACE2 bond with single-molecule precision.

Using the unfolding of ACE2 as a fingerprint pattern, we were able to bring together equilibrium bond dynamic studies using magnetic tweezers, high force rupture analysis carried out with an AFM, and molecular insights of the unfolding process obtained by SMD simulations. Combining these approaches, we were able to cover the whole physiologically relevant force range. We compared the force-stability of the SARS-CoV-1 and SARS-CoV-2 RBD to human receptor ACE2 and found higher force stability for the novel coronavirus throughout all force regimes. This is in line with previously published in silico and in vitro force spectroscopy studies that found a 20 to 40% difference in the force-stability of SARS-CoV-1 and SARS-CoV-2 RBD:ACE2 interactions (40, 41). In a binding environment that is permanently exposed to mechanical perturbations, this is an evolutionary advantage and hints at the value of force-spectroscopy to understand fitness advantages of newly evolving mutants.

Apart from bond stability, we analyzed the binding kinetics of the complex under force obtained by MT measurements, finding an order of magnitude higher bond lifetime of the SARS-COV-2 RBD construct compared to the same construct with the SARS-COV-1 RBD at their respective equilibrium forces. Extrapolating these lifetimes to zero force yields a more than 5-fold higher bond lifetime (∼115 s) for SARS-CoV-2 than for SARS-CoV-1 (∼20 s). We can quantitatively relate our results to studies that have reported equilibrium dissociation constants and rates for the ACE2 interactions with SARS-CoV-1 and SARS-CoV-2 using traditional binding assays. While the values reported in the literature vary significantly, likely due to the different experimental methods and sample immobilization strategies used, clear and consistent trends can be identified. The lifetimes of the bound complex determined in our assay correspond to rates of k_0,off_ ∼ 5.10^−2^ s^−1^ for SARS-CoV-1 and ∼8.7.10^−3^ s^−1^ for SARS-CoV-2, well within the ranges of reported k_sol,off_ values in literature (6, 24, 42-45) (for an overview see Supplementary Table 1). Our value for the off-rate of SARS-CoV-2 RBD bound to ACE2 is also in excellent agreement with the value of (8 ± 5).10^−3^ s^−1^ extrapolated from previous AFM force spectroscopy experiments (46). In contrast to the difference in off-rates between SARS-CoV-1 and SARS-CoV-2, the (bimolecular) on-rates reported in literature are similar for both SARS variants, in the range of ∼10^5^ M^-1^s^-1^. Consistently, our tethered ligand assay finds similar unimolecular on-rates on the order of ∼10 s^-1^. Even though the relative difference between both SARS viruses is similar, the absolute values are not comparable, as solution-based assays determine bulk on-rates, whereas our tethered ligand assay measures molecular on-rates. To relate the two quantities, an effective concentration c_eff_ can be introduced, such that k_sol,on_ = k_0,on_/c_eff_ (18, 47). This implies an effective concentration on the order of 0.1 mM. While the on-rate determined by the tethered ligand assay is not accurately reflecting the on-rate of both binding partners in solution, it mimics the naturally occurring effect of pre-formed interactions by a subset of the RBDs in the S trimer or by neighboring S trimers. This results in rapid formation of multivalent interactions between the virus and its host cell after an initial binding event, providing additional stability of the interaction. We estimate the concentration of S in vivo as ∼1 pM, based on 7.106 viral copies in ml sputum (8) and 100 S proteins per virus (1). This estimated bulk protein concentration in vivo is much lower than the dissociation constants reported, which are in the range K_d_ ∼ 1-100 nM for the SARS-CoV-2 RBD binding to ACE2 and 10-fold lower for SARS-CoV-1 (Supplementary Table 1), suggesting that multivalency might be critical for efficient viral binding. The rapid binding of RBDs held in proximity to ACE2 revealed by our assay might, therefore, be an important component of SARS-CoV-2 infections.

Our assay gives direct access to binding rates of receptor:ligand pairs held in spatial proximity. In addition, we can demonstrate and quantify blocking of the RBD:ACE2 bond. We anticipate that it will be a powerful tool to investigate the influence of mutations on interface stability and to assess the mode of action of potential therapeutic agents such as small molecules (48), neutralizing antibodies (42, 49), nanobodies (45, 50, 51), or designer proteins (52, 53) that interfere with S binding to ACE2. In particular, the tethered ligand assay could go beyond standard bulk assays and reveal heterogeneity, include avidity effects, the ability of direct displacement, and determine drug residence times, in addition to affinities. Investigating how mutations in the RBD affect force-stability might give valuable insights into reasons for fitness advantages of newly emerging variants and might even provide a tool to predict those advantages based on increased force-stability.

## MATERIALS AND METHODS

### Engineering of Recombinant Proteins, Experimental Procedures, and Data Analysis

SARS-CoV-1 and SARS-CoV-2 RBD:ACE2 tethered ligand constructs were designed based on available crystal structures PDB ID: 2ajf (23) and PDB ID: 6m0j (24). Please refer to the Supporting Information (SI) for details of how tagged protein constructs were expressed, purified and coupled to the surfaces for specific SMFS experiments. AFM and MT measurements were performed on custom-built instruments (26, 31) that were calibrated using the equipartition theorem to determine the spring constant of the AFM cantilever and the forces based on long DNA tethers in MT, respectively. The same SARS-CoV-1 and SARS-CoV-2 RBD:ACE2 constructs were investigated with both SMFS techniques under constant force-loading rate (AFM) and constant force (MT). Equilibrium measurements with MT and rupture experiments with an AFM were evaluated with custom MATLAB and Python scripts to deduce force stability and kinetics. SMD simulations were used to determine underlying molecular mechanisms. Details on the SMFS measurements, data analysis, and SMD simulations are given in the SI.

### Measurement conditions

Measurements of tethered ligand constructs in AFM and MT were performed in TRIS buffered saline (TBS - 25mM TRIS, 72mM NaCl, 1mM CaCl2 at pH 7.2 at RT). To test blocking, recombinant human ACE2 (Gln18-Ser740, C-terminal His-tag) from RayBiotech (∼3.8 µM in measurement buffer, # 230-30165-100 distributed by antibodies-online GmbH, # ABIN6952473) was flushed into the flowcell and shortly incubated before the measurement.

## Supporting information

Supplementary Infomation

## ACKNOWLEDGEMENTS

We thank David Dulin and Klaus Überla for helpful discussions and Nina Beier, Benedikt Böck, and Ellis Durner for help with initial experiments. This study was supported by German Research Foundation Projects 386143268 and 111166240, an EMBO long term fellowship to L.F.M. (ALTF 1047-2019), and the Physics Department of the LMU Munich.

## AUTHOR CONTRIBUTIONS

M.S.B., S.G., H.E.G., E.P., D.L., R.C.B., and J.L. designed research; M.S.B., S.G., and A.H. built instruments; M.S.B., S.G., A.H., and L.C.S. performed experiments; R.C.B. performed and analyzed simulations; M.S.B., P.L.N., E.P., T.N., L.F.M., and D.L. contributed new reagents/analytic tools; M.S.B., S.G., and A.H. analyzed experimental data; and M.S.B., S.G., H.E.G., and J.L. wrote the paper with input from all authors.

