## Supplementary Infomation for "A Tethered Ligand Assay to Probe SARS-CoV-2:ACE2 Interactions"

#### MATERIALS AND METHODS

All chemicals used, if not further specified, were supplied by Carl Roth (Karlsruhe, Germany) or Sigma-Aldrich (St. Louis, MO, USA).

##### Cloning and Protein Construct Design

Constructs for ACE2-linker-RBD of SARS-CoV-1 were designed in SnapGene Version 4.2.11 (GSL Biotech LLC, San Diego, CA, USA) based on a combination of the ACE2 sequence from Komatsu et al. (1) available from GenBank under accession number AB046569 and the SARS-CoV-1 sequence from Marra et al. (2) available from GenBank under accession number AY274119. The crystal structure by Li et al. (3) available from the Protein Data Bank (PDB ID: 2ajf) was used as a structural reference. The linker sequence and tag placement were adapted from Milles et al. (4). The linker sequence is a combination of two sequences available at the iGEM parts databank (accession numbers BBa_K404300, BBa_K243029). We used a similar approach to design the fusion protein with the sequence of the RBD of SARS-CoV-2 from the sequence published by Wu et al. (5) available from GenBank under accession number MN908947. A 6x histidine (His) tag was added for purification. In addition, tags for specific pulling in magnetic tweezers and the atomic force microscope were introduced: a triple glycine for sortase-catalyzed attachment on the N-terminus and a ybbR-tag, AviTag, and Fgɣ tag on the C-terminus. In summary, the basic construct is built up as follows: MGGG-ACE2-linker-RBD-6xHIS-ybbR-AviTag-Fgɣ. All protein sequences are provided in the full protein sequences paragraph.

The constructs were cloned using Gibson assembly from linear DNA fragments (GeneArt, ThermoFisher Scientific, Regensburg, Germany) containing the sequence of choice codon-optimized for expression in E. coli into a Thermo Scientific pT7CFE1-NHis-GST-CHA Vector (Product No. 88871). The control construct with a different sized linker and just ACE2 were obtained by blunt end cloning adding additional residues to the linker or deleting parts of the construct. Replication of DNA plasmids was obtained by transforming in DH5-Alpha Cells and running overnight cultures with 7 ml lysogeny broth with 50 µg/ml carbenicillin. Plasmids were harvested using a QIAprep® Spin Miniprep Kit (QIAGEN, Germantown, MD, USA, # 27106).

The used plasmids for force-spectroscopy measurements were deposited with and can be ordered from Addgene (www.addgene.org):

| Plasmid | AddgeneID |
|---|---|
| pT7CFE1-MGGG-ACE2-link33nm-SARS-CoV-1-RBD-HIS-ybbr-AviTag-Fgɣ | 174831 |
| pT7CFE1-MGGG-ACE2-link33nm-SARS-CoV-2-RBD-HIS-ybbr-AviTag-Fgɣ | 174832 |
| pT7CFE1-MGGG-ACE2-link42nm-SARS-CoV-1-RBD-HIS-ybbr-AviTag-Fgɣ | 174833 |
| pT7CFE1-MGGG-ACE2-HIS-ybbr-AviTag-Fgɣ | 174835 |

**In Vitro Protein Expression**

Expression was conducted according to the manual of 1-Step Human High-Yield Mini in vitro translation (IVT) kit (# 88891X) distributed by ThermoFisher Scientific (Pierce Biotechnology, Rockford, IL, USA). All components, except 5X dialysis buffer, were thawed on ice until completely thawed. 5X dialysis buffer was thawed for 15 minutes and 280 µl were diluted into 1120 µl nuclease-free water to obtain a 1X dialysis buffer. The dialysis device provided was placed into the dialysis buffer and kept at room temperature until it was filled with the expression mix.

For preparing the IVTT expression mix, 50 µl of the HeLa lysate was mixed with 10 µl of accessory proteins. After each pipetting step, the solution was gently mixed by stirring with the pipette. Then the HeLa lysate and accessory proteins mix was incubated for 10 minutes. Afterwards, 20 µl of the reaction mix was added. Then 8 µl of the specifically cloned DNA (0.5 µg/µl) was added. The reaction mix was then topped off with 12 µl of nuclease-free water to obtain a total of 100 µl. This mix was briefly centrifuged at 10,000 g for 2 minutes. A small white pellet appeared. The supernatant was filled into the dialysis device placed in the 1X dialysis buffer. The entire reaction was then incubated for 16 h at 30°C under constant shaking at 700 rpm. For incubation and shaking an ThermoMixer comfort 5355 (Eppendorf AG, Hamburg, Germany, # 5355) with a 2 ml insert was used. After 16 h the expression mix was removed and stored in a protein low binding reaction tube on ice until further use.

**Protein Purification**

Purification was conducted using HIS Mag Sepharose® Excel beads (Cytiva Europe GmbH, Freiburg, Germany, # 17371222) together with a MagRack™ 6 (Cytiva Europe GmbH, Freiburg, Germany, # 28948964) closely following the provided protocol. Bead slurry was mixed thoroughly by vortexing. 200 µl of homogenous beads were dispersed in a 1.5 ml protein low binding reaction tube. Afterwards the reaction tube was placed in the magnetic rack and the stock buffer was removed. Next, the beads were washed with 500 µl of HIS wash buffer (25 mM TRIS-HCl, 300 mM NaCl, 20 mM imidazole, 10% vol. glycerol, 0.25 % vol. Tween 20, pH 7.8). Expressed protein from IVTT was filled to 1000 µl with TRIS buffered saline (25 mM TRIS, 72 mM NaCl, 1 mM $CaCl_2$, pH 7.2) and mixed with freshly washed beads. The mix was incubated in a shaker for 1 h at room temperature. Subsequently, the reaction tube was placed in the magnetic rack and the liquid was removed. The beads were washed three times with wash buffer, keeping the total incubation time to less than 1 min. Remaining wash buffer was removed and 100 µl elution buffer (25 mM TRIS-HCl, 300 mM NaCl, 300 mM imidazole, 10% vol. glycerol, 0.25 % vol. Tween 20, pH 7.8) was added to wash protein off the beads. The bead elution buffer mix was then incubated for one minute with occasional gentle vortexing. Afterward, the reaction tube was placed in the magnetic rack again to remove the eluted protein. This step was repeated for a second and third elution step. The buffer of the eluted protein was exchanged to TRIS buffered saline (TBS - 25mM TRIS, 72mM NaCl, 1mM $CaCl_2$ at pH 7.2) in 0.5 ml 40k Zeba spin columns distributed by ThermoFisher Scientific (Pierce Biotechnology, Rockford, IL, USA, # 87767) or 0.5 ml 50k Amicon Centrifugal Filters (Merck KGaA, Darmstadt, Germany, #UFC5050BK). Concentrations were determined photospectrometrically with a NanoDrop and aliquots were frozen in liquid nitrogen.

**Atomic Force Microscopy Setup**

The AFM force spectroscopy datasets were collected on a custom-built AFM based on an MFP3D controller (Asylum Research, Santa Barbara, CA, USA). The AFM head was kept stationary, while the sample stage was moved by an xyz-movable piezo-driven sample stage (P-313.30D - P-313 PicoCube® XY(Z)-Piezoscanner - Physik Instrumente PI GmbH & Co KG, Karlsruhe, Germany) together with a high-precision xy-nanopositioner (P-621.2CD - Physik Instrumente PI GmbH & Co KG, Karlsruhe, Germany). Unfolding traces were was recorded by (i) approaching the functionalized sample surface onto the functionalized cantilever until the cantilever is indented with 180 pN into the surface, allowing the linkage between ClfA:Fgɣ to form; (ii) retraction of the cantilever with 800 nm/s while recording the deflection of the cantilever to obtain a force distance curve of the mechanical response of the protein probed; (iii) after the surface moved 500 nm in z direction (assuming a complete unfolding) a new position is set by the xy-stage, moving the sample surface horizontally by steps of 100 nm in a spiral pattern and starting a new acquisition process in step (i). This process is operated by a software programmed in IgorPro6 (Wavemetrics, Portland, OR, USA) and the obtain unfolding curves saved in an hdf5 file.

For calibration the Inverse Optical Cantilever Sensitivity (InvOLS) was obtained by 25 hard indentation curves allowing to correlate the movement of the cantilever with the voltage signal recorded on a quadrant photodetector. The spring constants of the cantilevers were calibrated based on thermal fluctuations using the equipartition theorem method (6, 7) resulting in spring constants around 100 pN/nm. The spring constant per measurement are listed below. To be able to directly compare force values recorded with different cantilevers, the ACE2 fingerprint pattern was used to normalize the force histograms.

In the course of a measurement around 50,000 - 400,000 force-distance traces were recorded. Most force-distance traces didn't show any interactions and only a fraction showed specific single-molecule unfoldings.

List of spring constants (k) of cantilevers used for individual measurements used for comparing forces and contour length increments:

SARS-CoV-2 measurement - Figure 2A (exemplary curve), C and Supplementary Figure S1

k = 101.4 pN/nm

SARS-CoV-1 short linker (33 nm) measurement - Figure 2B

k = 108.0 pN/nm

SARS-CoV-1 long linker (42 nm) measurement - Figure 2B top only

k = 109.5 pN/nm

Ectodomain of ACE2 - Supplementary Figure S3

k = 96.0 pN/nm

**AFM surface and cantilever preparation**

Cantilevers and sample surfaces were both silanzied for further functionalization steps. Surface attachment and linkage was obtained by 5,000 Da heterobifunctional NHS-PEG-Maleimide spacers (Rapp Polymere, Tübingen, Germany, # 135000-65-35). Specific protein attachment was achieved using a sortase-mediated reaction on the sample surface and an sfp-mediated reaction for the AFM cantilever ensuring a well-defined pulling geometry.

The cantilevers (BioLever mini, BL-AC40TS) were oxidized in an UV ozone cleaner (UVOH 150 LAB; FHR Anlagenbau GmbH, Ottendorf-Okrilla, Germany) and after that silanized for 2 min in 50% (vol/vol) ethanol and (3-aminopropyl)dimethylethoxysilane (abcr GmbH, Karlsruhe, Germany, AB146193, CAS 18306-79-1). To rinse off the residual silane cantilevers were stirred in 2-Propanol (IPA) and in MilliQ. After that the cantilevers are dried at 80°C for 30 min. The heterobifunctional PEG spacers are solved in 100 mM HEPES (pH 7.4) to 50 mM. The cantilevers incubated in droplets of that mixture for 1 h. After a rinsing step the cantilevers are incubated in 20 mM Coenzyme A (CoA) (# 234101-100MG,

Calbiochem distributed by Sigma-Aldrich) dissolved in coupling buffer (50 mM Disodium phosphate buffer, 50 mM NaCl, 10 mM EDTA, pH 7.2). for 1 h. Droplets with a mixture of 2 µM, 100 mM $MgCl_2$ and 60 µM ClfA (in TBS buffer) are prepared to attach the ybbR tag of ClfA specifically to CoA. The cantilever are incubated in these droplets for at least 1.5 h. To prepare the cantilevers for the measurement they are rinsed and store in TBS.
To clean the glass surfaces for silanized they are sonicated in 50% (vol/vol) IPA in MilliQ for 15 min. To prime the surfaces for silanization they are incubated for 30 min in a solution of 50% (vol/vol) hydrogen peroxide (30%) and sulfuric acid. To wash off the residual solution the surfaces are washed in MilliQ and dried under a constant stream of N2. For the actual silanization step the surfaces are incubated in 1.8% (vol/vol) ethanol and (3-aminopropyl)dimethylethoxysilane (abcr GmbH, Karlsruhe, Germany, AB146193, CAS 18306-79-1). Afterwards the surfaces are washed with IPA and MilliQ and dried at 80°C for 45 min. To minimize sample volumes for the following incubations silicone incubation wells (CultureWell reusable gaskets, Grace Bio-Labs, Bend, OR, USA, # 103250) are placed centered on the surfaces. Then heterobifunctional PEG spacers are solved in 100 mM HEPES (pH 7.4) to 50 mM and applied to the wells for 1 h. After that the surfaces are rinsed with MilliQ and 5 mM Cys-LPETGG in coupling buffer (sodium phosphate, pH 7.2) is pipetted in the wells and incubated for 2 h. After washing the wells the purified tethered ligand protein is applied in solution together with 1 µM sortase and 10 mM CaCl2. After incubation the well is removed and the surface rinsed with 10 ml TBS.
As a control surface to validate the functionality of the cantilever and of the surfaces another surface with a GGG-ddfln4-Fgɣ harboring an Fgɣ tag is prepared analog to the actual sample surface.

**AFM data analysis**
Data analysis was carried out in custom Python 2.7 (Python Software Foundation, Wilmington, DE, USA) scripts (8-10) and Python 2.7-based Jupyter notebooks (11).
The rupture forces were detected by a peak detection highlighting drops above the baseline noise level of Total Variation Denoised (TVD) force-distance traces. A linear slope in force vs. time was used to determine the loading rate, taking into account 4 nm before the rupture event. Rupture forces of respective domains in the unfolding pattern were binned to histograms and fitted with the Bell-Evans model yielding the most probable rupture force (12).
All curves showing the characteristic unfolding pattern of the tethered ligand protein were aligned (accounting for the inhomogeneity in PEG lenghts) and assembled to heatmaps to visualize the recurring, characteristic unfolding pattern. The heatmaps contain the raw unfolding curves in force-distance space binned in 90 bins in both x- and y-axis between -10 pN to 60 pN and -10 nm to 300 nm.
To transform force-extension data into contour length space (13) a three-regime model by Livadaru et. al (14) assuming a stiff element of b = 0.11 nm and bond angle γ = 41° was used. A Gaussian kernel density estimate, with a bandwidth of 4 nm, was applied to the gained contour length data to obtain density curves of each trace. These curves were aligned in contour length space using the following process previously described by Baumann et al. (15): “the full set of transformed force-distance curves is aligned to a random curve from this data set according to least residual in cross-correlation. This process results in a first superposition which is used as a template in a second iteration of this process. Again, all contour-length transformed curves are aligned to a template curve but this time to the one formed by the first iteration. This two-step approach diminishes biasing effects given by the choice of the random curve used for initial alignment. Contour lengths of the individual domains are determined by a Gaussian fit of each determined peak and subtraction of the respective fitted means.”

**Molecular dynamic simulation**
To examine the stability of the protein complex under mechanical load, we carried out steered molecular dynamics simulations employing NAMD 3 (16). Simulations were prepared using VMD (17) and its QwikMD (18) molecular dynamics interface. The structure of the

complexes were prepared following established protocols (19). For the SARS-CoV-1 RBD:ACE2 complex, the structure had been solved by means of X-ray crystallography at 2.90 Å resolution and is available at the protein data bank (PDB ID: 2ajf) (3). For the SARS-CoV-2 RBD:ACE2 complex, the structure had been solved by means of X-ray crystallography at 2.90 Å resolution and is available at the protein data bank (PDB ID: 6moj) (20) SARS-CoV-1 RBD or SARS-CoV-2 RBD and the ectodomain of human ACE2 were joined by flexible polypeptide linkers. The structure of the complexes with the linkers was obtained using MODELLER (21) and fitted with VMD (REF). Disulfide bonds were included following the literature information (22). All structures were subjected to 10 ns of equilibrium MD, to ensure conformational stability.

Employing advanced run options of QwikMD, structural models were solvated and the net charge of the proteins were neutralized using a 75 mM salt concentration of sodium chloride, which were randomly arranged in the solvent. The overall number of atoms included in MD simulations varied from approximately 200,000 in the RDB:ACE2 systems with no linker, to nearly 4,000,000 in the systems RDB:ACE2 connected by flexible polypeptide linkers. All simulations were performed employing the NAMD molecular dynamics package (16), taking advantage of NVIDIA DGX-A100-based cluster nodes at Auburn University. The CHARMM force field (23, 24) along with the TIP3 water model (25) was used to describe all systems. The simulations were performed assuming periodic boundary conditions in the NpT ensemble with temperature maintained at 300 K using Langevin dynamics for pressure, kept at 1 bar, and temperature coupling. A distance cut-off of 12.0 Å was applied to short-range, non-bonded interactions, whereas long-range electrostatic interactions were treated using the particle-mesh Ewald (PME) (26) method. The equations of motion were integrated using the r-RESPA multiple time step scheme (27) to update the van der Waals interactions every two steps and electrostatic interactions every four steps. The time step of integration was chosen to be 4 fs for all production simulations performed, and 2 fs for all equilibration runs. For the 4 fs simulations, hydrogen mass repartitioning was done using psfgen in VMD. Before the MD simulations all the systems were submitted to an energy minimization protocol for 5,000 steps.

MD simulations with position restraints in the protein backbone atoms were performed for 1.0 ns and served to pre-equilibrate systems before the 10 ns equilibrium MD runs, which served to evaluate structural model stability. During the 1.0 ns pre-equilibration the initial temperature was set to zero and was constantly increased by 1 K every 1,000 MD steps until the desired temperature (300 K) was reached. SMD simulations (28) were performed using a constant velocity stretching (SMD-CV protocol), employing three different pulling speeds: 12.5, 2.5, 0.25 and 0.025 Å/ns. In our *in silico* SMFS approach, many replicas of simulations are performed (at least 24 per system). The simulation replicas, used in all the plots in this manuscript, were performed with constant pulling speed of 2.5 Å/ns. Values for force over the pulling spring were saved every 50 steps. The spring constant of the pulling spring was set to 5.0 kcal/mol/Å, while the holding spring had a constant of 100 kcal/mol/Å. In all simulations, totaling over 300 SMD simulations, SMD was employed by harmonically restraining the position of a terminal amino acid residue, and moving a second restraint point, at a terminal amino acid residue of the other domain, with constant velocity in the +z direction. The procedure is equivalent to attaching one end of a harmonic spring to the end of a domain and pulling on the other end of the spring. The force applied to the harmonic spring is then monitored during the time of the molecular dynamics simulation. The pulling point was moved with constant velocity along the z-axis and due to the single anchoring point and the single pulling point the system is quickly aligned along the z-axis. Owing to the flexibility of the linkers between the RDB:ACE2 and fingerprint domains, this approach reproduces the experimental set-up.

Analyses of MD trajectories were carried out employing VMD (17) and its plug-ins, as well as in-house python-based Jupyter notebooks (11). Secondary structures were assigned using the Timeline plug-in, which employs STRIDE criteria. Force propagation profiles (29) were analysed using generalized cross correlation-based network analysis (30) A network was defined as a set of nodes, all α-carbons, with connecting edges. Edges connect pairs of nodes if corresponding monomers are in contact, and 2 non-consecutive monomers are said to be in contact if they fulfill a proximity criterion, namely any heavy atoms (non-hydrogen) from the 2 monomers are within 4.5Å of each other for at least 75% of the frames analyzed.

**Magnetic Tweezers Instrument**

Measurements were performed on a custom MT setup described previously (31, 32). In the setup, molecules are tethered in a flow cell (FC; see next section); mounted above the FC is a pair of permanent magnets (5×5×5 mm3 each; W-05-N50-G, Supermagnete, Gottmadingen, Germany) in vertical configuration (33). The distance between magnets and FC is controlled by a DC-motor (M-126.PD2, Physik Instrumente PI GmbH & Co KG, Karlsruhe, Germany) and the FC is illuminated by an LED (69647, Lumitronix LED Technik GmbH, Germany). Using a 40x oil immersion objective (UPLFLN 40x, Olympus, Japan) and a CMOS sensor camera with 5120 x 5120 pixels (5120 x 5120 pixels, CP80-25-M-72, Optronis, Kehl, Germany) a field of view of approximately 680 × 680 $\mu m^2$ is imaged at a frame rate of 72 Hz. Images are transferred to a frame grabber (microEnable 5 ironman VQ8-CXP6D, Silicon Software, Mannheim, Germany) and analyzed with an open-source tracking software (34, 35). The tracking accuracy of our setup was determined to be ≈ 0.6 nm in (x, y) and ≈ 1.5 nm in z direction, as determined by tracking non-magnetic polystyrene beads, after baking them onto the flow cell surface. For creating the look-up table required for tracking the bead positions in z, the objective is mounted on a piezo stage (Pifoc P-726.1CD, Physik Instrumente PI GmbH & Co KG, Karlsruhe, Germany). Force calibration was performed as described (36) by analysis of the fluctuations of long DNA tethers. Importantly, for the small extension changes on the length scales of our protein tethers, the force stays constant to very good approximation (to better than 10−4 relative change). The largest source of force uncertainty is due to bead-to-bead variation, which is on the order of ≤ 10% for the beads used in this study (33, 37).

**Flowcell Preparation and Magnetic Tweezers Measurements**

Flowcells (FCs) were prepared as described previously (31). High precision microscope cover glasses (24 mm x 60 mm x 0.17 mm, Carl Roth) were amino-silanized for further functionalization (equal to AFM surface preparation). They were coated with sulfo-SMCC (38) (sulfosuccinimidyl 4-(N-maleimidomethyl)cyclohexane-1-carboxylate; sulfo-SMCC, ThermoFisher Scientific, Pierce Biotechnology, Rockford, IL, USA, # 22322). For this purpose, 180 µl sulfo-SMCC (10 mM in 50 mM Hepes buffer, pH 7.4) was applied to one amino-silanized slide that was sandwiched with another slide and incubated for 45 min. Unbound sulfo-SMCC was removed by rinsing with MilliQ. Next, elastin-like polypeptide (ELP) linkers (39) with a sortase motif at their C-terminus were coupled to the maleimide of the sulfo-SMCC via a single cysteine at their N-terminus, by sandwiching two slides with 100 µl ELP linkers (in 50 mM Disodium phosphate buffer with 50mM NaCl and 10mM EDTA, pH 7.2) and incubating them for 60 min. Subsequently, after further MilliQ rinsing to remove unbound ELP linkers, free sulfo-SMCC was neutralized with free cysteine (10 mM in 50 mM Disodium phosphate buffer with 50 mM NaCl and 10 mM EDTA, pH 7.3). 1 µm diameter polystyrene beads dissolved in ethanol were applied to the glass slides. After the ethanol evaporated, beads were baked onto the glass surface to serve as reference beads during the measurement. FCs were assembled from an ELP-functionalized bottom slide and an unfunctionalized high-precision microscope cover glass slide with two holes (inlet and outlet) on either side serving as top slide. Both slides were separated by a layer of parafilm (Pechiney Plastic Packaging Inc., Chicago, IL, USA), which was cut out to form a 50 µl channel. FCs were incubated with 1% (v/v) casein solution (# C4765-10ML, Sigma-Aldrich) for 2 h and flushed with 1 ml buffer (25 mM TRIS, 72 mM NaCl, 1 mM CaCl2, pH 7.2 at RT).
CoA-biotin (# S9351 discontinued, New England Biolabs, Frankfurt am Main, Germany) was coupled to the ybbR-tag at the C-terminus of the fusion protein constructs in a 90 - 120 min bulk reaction in the presence of 4 µM sfp phosphopantetheinyl transferase (40) and 100 mM $MgCl_2$ at room temperature (≈ 22°C). Proteins were diluted to a final concentration of about 50 nM in 25 mM TRIS, 72 mM NaCl, 1 mM $CaCl_2$, pH 7.2 at RT. To couple the N-terminus of the fusion proteins carrying three glycines to the C-terminal LPETGG motif of the ELP-linkers, 100 µl of the protein mix was flushed into the FC and incubated for 24 min in the

presence of 1.3 µM evolved pentamutant sortase A from Staphylococcus aureus (41, 42). Unbound proteins were flushed out with 1 ml measurement buffer (25 mM TRIS, 72 mM NaCl, 1 mM CaCl2, 0.1% (v/v) Tween-20, pH 7.2 at RT). Finally, commercially available streptavidin-coated paramagnetic beads (Dynabeads™ M-270 Streptavidin, Invitrogen, Life Technologies, Carlsbad, CA, USA) were added into the FC and incubated for 30 s before flushing out unbound beads with 1 ml measurement buffer. Receptor-ligand binding and unbinding under force was systematically investigated by subjecting the protein tethers to (2 - 30) min long plateaus of constant force, which was gradually increased in steps of 0.2 or 0.3 pN. All measurements were conducted at room temperature.
For blocking measurements, recombinant human ACE2 (Gln18-Ser740, C-terminal His-tag) from RayBiotech (Peachtree Corners, GA, USA, # 230-30165-100 distributed by antibodies-online GmbH, Aachen, Germany, # ABIN6952473) was dissolved in measurement buffer for a final concentration of ~3.8 µM. Dissolved ACE2 was spinned down in a tabletop centrifuge at 4°C, 14,000 rcf for 5 min to avoid introduction of larger particles into the FC that could influence video tracking. 80 µl ACE2 were flushed into the FC and shortly incubated before applying 7 pN to force dissociation of the tethered ligand construct and allow the free ACE2 to bind. Afterwards, a measurement was conducted in the presence of free ACE2.

**Data Analysis of MT traces**

MT traces for thermodynamic bond analysis were selected on the basis of the characteristic ACE2 two-step unfolding pattern above 25 pN, conducted at the end of each experiment. For each trace, (x,y)-fluctuations were also checked to avoid inclusion of tethers that exhibit inter-bead or bead-surface interactions, which would also cause changes in x or y. Non-magnetic reference beads were tracked simultaneously with magnetic beads and reference traces were subtracted for all measurements to correct for drift. Extension time traces were subjected to a 5-frame moving average smoothing to reduce noise. All analyses were performed with custom scripts in MATLAB.

**Full protein sequences**

**pT7CFE1-MGGG-ACE2-link33nm-SARS-CoV-1-RBD-HIS-ybbr- AviTag-Fgɣ**

MGGGSSSTIEEQAKTFLDKFNHEAEDLFYQSSLASWNYNTNITEENVQNMNNAGDK
WSAFLKEQSTLAQMYPLQEIQNLTVKLQLQALQQNGSSVLSEDKSKRLNTILNTMST
IYSTGKVCNPDNPQECLLLEPGLNEIMANSLDYNERLWAWESWRSEVGKQLRPLYE
EYVVLKNEMARANHYEDYGDYWRGDYEVNGVDGYDYSRGQLIEDVEHTFEEIKPL
YEHLHAYVRAKLMNAYPSYISPIGCLPAHLLGDMWGRFWTNLYSLTVPFGQKPNIDV
TDAMVDQAWDAQRIFKEAEKFFVSVGLPNMTQGFWENSMLTDPGNVQKAVCHPT
AWDLGKGDFRILMCTKVTMDDFLTAHHEMGHIQYDMAYAAQPFLLRNGANEGFHE
AVGEIMSLSAATPKHLKSIGLLSPDFQEDNETEINFLLKQALTIVGTLPFTYMLEKWR
WMVFKGEIPKDQWMKKWWEMKREIVGVVEPVPHDETYCDPASLFHVSNDYSFIRY
YTRTLYQFQFQEALCQAAKHEGPLHKCDISNSTEAGQKLFNMLRLGKSEPWTLALE
NVVGAKNMNVRPLLNYFEPLFTWLKDQNKNSFVGWSTDWSPYADGATSGGGGSA
GGSGSGSSGGSSGASGTGTAGGTGSGSGTGSGGGSGGGSEGGGSEGGGSEGG
GSEGGGSEGGGSGGGSESGGSSARVVPSGDVVRFPNITNLCPFGEVFNATKFPSV
YAWERKKISNCVADYSVLYNSTFFSTFKCYGVSATKLNDLCFSNVYADSFVVKGDD
VRQIAPGQTGVIADYNYKLPDDFMGCVLAWNTRNIDATSTGNYNYKYRYLRHGKLR
PFERDISNVPFSPDGKPCTPPALNCYWPLNDYGFYTTTGIGYQPYRVVVLSFELLNA
PATVCGPKLSTDLIKNQCVNFSGHHHHHHTDSLEFIASKLAASGLNDIFEAQKIEWHE
GSGEGQQHHLGGAKQAGDV*

**pT7CFE1-MGGG-ACE2-link33nm-SARS-CoV-2-RBD-HIS-ybbr-AviTag-Fgɣ**

MGGGSSSTIEEQAKTFLDKFNHEAEDLFYQSSLASWNYNTNITEENVQNMNNAGDK
WSAFLKEQSTLAQMYPLQEIQNLTVKLQLQALQQNGSSVLSEDKSKRLNTILNTMST
IYSTGKVCNPDNPQECLLLEPGLNEIMANSLDYNERLWAWESWRSEVGKQLRPLYE
EYVVLKNEMARANHYEDYGDYWRGDYEVNGVDGYDYSRGQLIEDVEHTFEEIKPL
YEHLHAYVRAKLMNAYPSYISPIGCLPAHLLGDMWGRFWTNLYSLTVPFGQKPNIDV
TDAMVDQAWDAQRIFKEAEKFFVSVGLPNMTQGFWENSMLTDPGNVQKAVCHPT
AWDLGKGDFRILMCTKVTMDDFLTAHHEMGHIQYDMAYAAQPFLLRNGANEGFHE
AVGEIMSLSAATPKHLKSIGLLSPDFQEDNETEINFLLKQALTIVGTLPFTYMLEKWR
WMVFKGEIPKDQWMKKWWEMKREIVGVVEPVPHDETYCDPASLFHVSNDYSFIRY
YTRTLYQFQFQEALCQAAKHEGPLHKCDISNSTEAGQKLFNMLRLGKSEPWTLALE
NVVGAKNMNVRPLLNYFEPLFTWLKDQNKNSFVGWSTDWSPYADGATSGGGGSA
GGSGSGSSGGSSGASGTGTAGGTGSGSGTGSGGGSGGGSEGGGSEGGGSEGG
GSEGGGSEGGGSGGGSESGGSSASNFRVQPTESIVRFPNITNLCPFGEVFNATRF
ASVYAWNRKRISNCVADYSVLYNSASFSTFKCYGVSPTKLNDLCFTNVYADSFVIRG
DEVRQIAPGQTGKIADYNYKLPDDFTGCVIAWNSNNLDSKVGGNYNYLYRLFRKSN
LKPFERDISTEIYQAGSTPCNGVEGFNCYFPLQSYGFQPTNGVGYQPYRVVVLSFEL
LHAPATVCGPKKSTNLVKNSGHHHHHHTDSLEFIASKLAASGLNDIFEAQKIEWHEG
SGEGQQHHLGGAKQAGDV*

**pT7CFE1-MGGG-ACE2-link42nm-SARS-CoV-1-RBD-HIS-ybbr-AviTag-Fgɣ**

MGGGSSSTIEEQAKTFLDKFNHEAEDLFYQSSLASWNYNTNITEENVQNMNNAGDK
WSAFLKEQSTLAQMYPLQEIQNLTVKLQLQALQQNGSSVLSEDKSKRLNTILNTMST
IYSTGKVCNPDNPQECLLLEPGLNEIMANSLDYNERLWAWESWRSEVGKQLRPLYE
EYVVLKNEMARANHYEDYGDYWRGDYEVNGVDGYDYSRGQLIEDVEHTFEEIKPL
YEHLHAYVRAKLMNAYPSYISPIGCLPAHLLGDMWGRFWTNLYSLTVPFGQKPNIDV
TDAMVDQAWDAQRIFKEAEKFFVSVGLPNMTQGFWENSMLTDPGNVQKAVCHPT
AWDLGKGDFRILMCTKVTMDDFLTAHHEMGHIQYDMAYAAQPFLLRNGANEGFHE
AVGEIMSLSAATPKHLKSIGLLSPDFQEDNETEINFLLKQALTIVGTLPFTYMLEKWR
WMVFKGEIPKDQWMKKWWEMKREIVGVVEPVPHDETYCDPASLFHVSNDYSFIRY
YTRTLYQFQFQEALCQAAKHEGPLHKCDISNSTEAGQKLFNMLRLGKSEPWTLALE
NVVGAKNMNVRPLLNYFEPLFTWLKDQNKNSFVGWSTDWSPYADGSEGGGSEGG
GSEGGGSEGGGSEGGGSGGGGATSGGGGSAGGSGSGSSGGSSGASGTGTAGG
TGSGSGTGSGGGSGGGSEGGGSEGGGSEGGGSEGGGSEGGGSGGGSESGGSS

ARVVPSGDVVRFPNITNLCPFGEVFNATKFPSVYAWERKKISNCVADYSVLYNSTFF
STFKCYGVSATKLNDLCFSNVYADSFVVKGDDVRQIAPGQTGVIADYNYKLPDDFM
GCVLAWNTRNIDATSTGNYNYKYRYLRHGKLRPFERDISNVPFSPDGKPCTPPALN
CYWPLNDYGFYTTTGIGYQPYRVVVLSFELLNAPATVCGPKLSTDLIKNQCVNFSGH
HHHHHTDSLEFIASKLAASGLNDIFEAQKIEWHEGSGEGQQHHLGGAKQAGDV*

**pT7CFE1-MGGG-ACE2-HIS-ybbr-AviTag-Fgɣ**

MGGGSSSTIEEQAKTFLDKFNHEAEDLFYQSSLASWNYNTNITEENVQNMNNAGDK
WSAFLKEQSTLAQMYPLQEIQNLTVKLQLQALQQNGSSVLSEDKSKRLNTILNTMST
IYSTGKVCNPDNPQECLLLEPGLNEIMANSLDYNERLWAWESWRSEVGKQLRPLYE
EYVVLKNEMARANHYEDYGDYWRGDYEVNGVDGYDYSRGQLIEDVEHTFEEIKPL
YEHLHAYVRAKLMNAYPSYISPIGCLPAHLLGDMWGRFWTNLYSLTVPFGQKPNIDV
TDAMVDQAWDAQRIFKEAEKFFVSVGLPNMTQGFWENSMLTDPGNVQKAVCHPT
AWDLGKGDFRILMCTKVTMDDFLTAHHEMGHIQYDMAYAAQPFLLRNGANEGFHE
AVGEIMSLSAATPKHLKSIGLLSPDFQEDNETEINFLLKQALTIVGTLPFTYMLEKWR
WMVFKGEIPKDQWMKKWWEMKREIVGVVEPVPHDETYCDPASLFHVSNDYSFIRY
YTRTLYQFQFQEALCQAAKHEGPLHKCDISNSTEAGQKLFNMLRLGKSEPWTLALE
NVVGAKNMNVRPLLNYFEPLFTWLKDQNKNSFVGWSTDWSPYADGSGHHHHHHT
DSLEFIASKLAASGLNDIFEAQKIEWHEGSGEGQQHHLGGAKQAGDV*

**pET28a_ClfA_N2N3-HIS-ybbr-LPETGG**

MATAPVAGTDITNQLTNVTVGIDSGTTVYPHQAGYVKLNYGFSVPNSAVKGDTFKITVPK
ELNLNGVTSTAKVPPIMAGDQVLANGVIDSDGNVIYTFTDYVNTKDDVKATLTMPAYIDPE
NVKKTGNVTLATGIGSTTANKTVLVDYEKYGKFYNLSIKGTIDQIDKTNNTYRQTIYVNPS
GDNVIAPVLTGNLKPNTDSNALIDQQNTSIKVYKVDNAADLSESYFVNPENFEDVTNSVNI
TFPNPNQYKVEFNTPDDQITTPYIVVVNGHIDPNSKGDLALRSTLYGYNSNIIWRSMSWD
NEVAFNNGSGSGDGIDKPVVPEQPSGHHHHHHGSDSLEFIASKLASLPETGG*

**pET28a_MGGG-ybbr-HIS-ddFLN4(C18S)-Fgɣ**

MGGGDSLEFIASKLAHHHHHHGSADPEKSYAEGPGLDGGESFQPSKFKIHAVDPD
GVHRTDGGDGFVVTIEGPAPVDPVMVDNGDGTYDVEFEPKEAGDYVINLTLDGDN
VNGFPKTVTVKPAPGSGSGSGSGEGQQHHLGGAKQAGDV*

### TABLES

| Study | ACE2 binding to SARS-CoV-1 RBD | ACE2 binding to SARS-CoV-2 RBD | Method and Comments |
|---|---|---|---|
| Lan *et al.* (20) | $K_d$ = 31 nM<br>$k_{sol,off} = 4.3\times10^{-2}$ s$^{-1}$<br>$k_{\mathrm{sol,on}} = 1.4\times10^{6}$ s$^{-1}$M$^{-1}$ | $K_{\mathrm{d}}$ = 4.7 nM<br>$k_{sol,off} = 6.5\times10^{-3}$ s$^{-1}$<br>$k_{\mathrm{sol,on}} = 1.4\times10^{6}$ s$^{-1}$M$^{-1}$ | Surface-plasmon resonance |
| Shang *et al.* (43) | $K_d$ =185 nM<br>$k_{sol,off} = 3.7\times10^{-2}$ s$^{-1}$<br>$k_{\mathrm{sol,on}} = 2.0\times10^{5}$ s$^{-1}$M$^{-1}$ | $K_d$ =44.2 nM<br>$k_{sol,off} = 7.8\times10^{-3}$ s$^{-1}$<br>$k_{\mathrm{sol,on}} = 1.75\times10^{5}$ s$^{-1}$M$^{-1}$ | Surface-plasmon resonance |
| Starr *et al.* (44) | $K_d$ =0.12 nM | $K_d$ =0.039 nM | Yeast display screen |
| Walls *et al.* (45) | $K_{\mathrm{d}}$ = 5.0 ± 0.1 nM<br>$k_{sol,off}$ = (8.7 ± 5.1) $\times10^{-4}$ s$^{-1}$<br>$k_{\mathrm{sol,on}}$ = (1.7 ± 0.7) $\times10^{5}$ s$^{-1}$M$^{-1}$ | $K_{\mathrm{d}}$ = 1.2 ± 0.1 nM<br>$k_{sol,off}$ = (1.7 ± 0.8) $\times10^{-4}$ s$^{-1}$<br>$k_{\mathrm{sol,on}}$ = (2.3 ± 1.4) $\times10^{5}$ s$^{-1}$M$^{-1}$ | Bio-layer interferometry; uses S protein for both variants |
| Wang *et al.* (46) | $K_d$ = 408 ± 11 nM<br>$k_{\mathrm{sol,off}}$ = (1.9 ± 0.4) $\times10^{-3}$ s$^{-1}$<br>$k_{\mathrm{sol,on}}$ = (2.9 ± 0.2) $\times10^{5}$ s$^{-1}$M$^{-1}$ | $K_d$ = 95 ± 7 nM<br>$k_{sol,off}$ = (3.8 ± 0.2) $\times10^{-3}$ s$^{-1}$<br>$k_{\mathrm{sol,on}}$ = (4.0 ± 0.2)$\times10^{4}$ s$^{-1}$M$^{-1}$ | Surface-plasmon resonance; uses S1 domain for SARS-CoV-2 |
| Wrapp *et al.* (47) | $K_{\mathrm{d}}$ = 325 nM<br>$k_{sol,off} = 112\times10^{-3}$ s$^{-1}$<br>$k_{\mathrm{sol,on}} = 3.62\times10^{5}$ s$^{-1}$M$^{-1}$ | $K_{\mathrm{d}}$ = 14.7 nM<br>$k_{sol,off} = 2.76\times10^{-3}$ s$^{-1}$<br>$k_{\mathrm{sol,on}}$ = 1.88 $\times10^{5}$ s$^{-1}$M$^{-1}$ | Surface-plasmon resonance; uses ectodomain for both variants |

**Supplementary Table 1. Equilibrium binding data for ACE2 binding to SARS-CoV-1 or SARS-CoV-2 RBD or S1 proteins.** Studies for both ACE2 binding to RBD constructs and to the S protein are included; Wrapp et al. find $K_d$ = 14.7 nM for ACE2 binding to SARS-CoV-2 S and $K_d$ = 34.6 nM for ACE2 binding to SARS-CoV-2 RBD, indicating similar affinities. Similarly, Yang et al. observe similar binding constants and mechanical stabilities for ACE2 binding to either the RBD or S using AFM force spectroscopy (48).

**FIGURES**

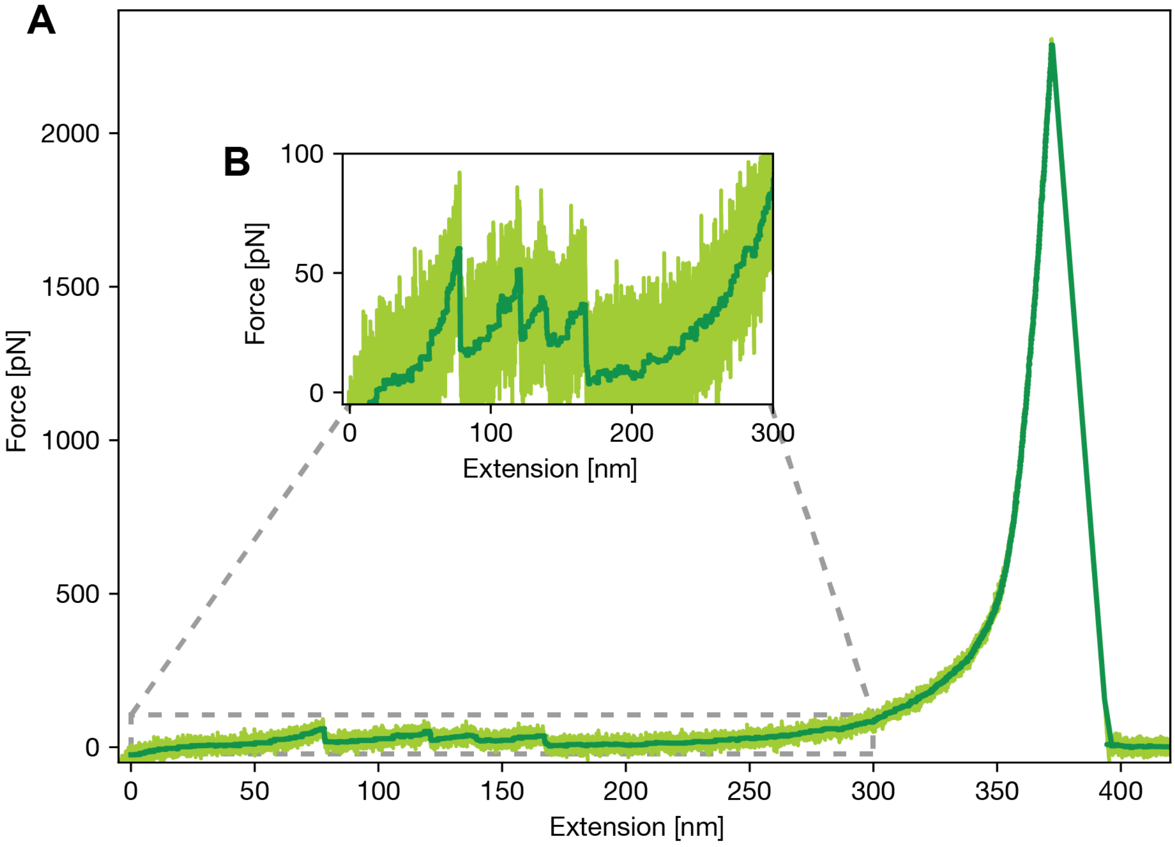


**Supplementary Fig. 1: Full force extension curve of the tethered ligand protein construct including the RBD of SARS-CoV-2 and ACE2. A** Complete force extension curve showing the entire unfolding until the final rupture of the ClfA:Fgɣ linkage. The full curve shows 4 peaks at lower forces (< 100 pN) and one final rupture at high forces (> 1000 pN). **B** Inset shows the extension range between 0 and 300 nm showing the low force unfolding events attributed to the tethered ligand protein. The first low force peak can be identified as the interface unbinding between SARS-CoV-2 RBD and ACE2 together with a partial unfolding of the RBD. This peak is followed by a trident shaped, three peak pattern that can be assigned to the unfolding of ACE2, see main text.

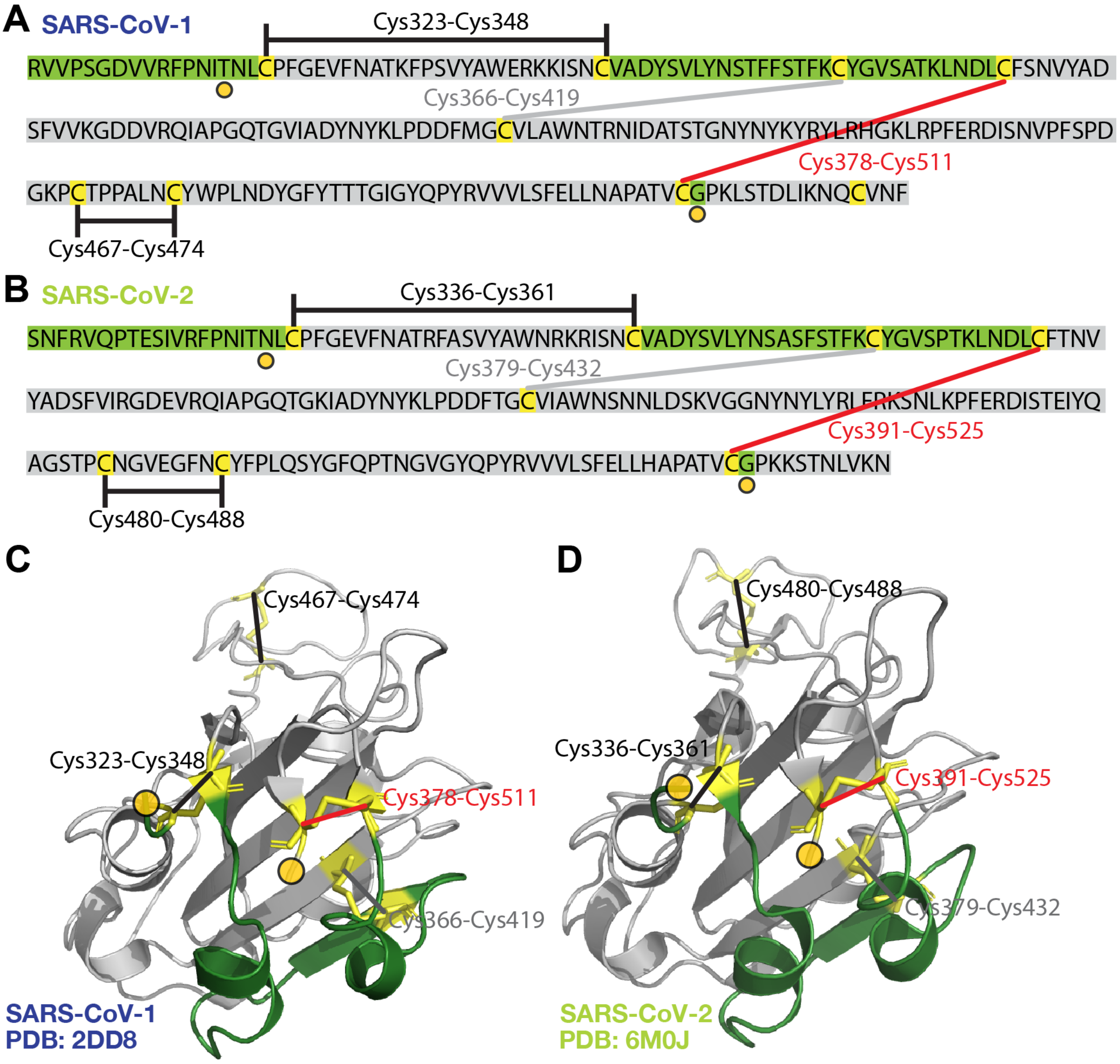


**Supplementary Fig. 2** Disulfide bridges shield large parts of the SARS-CoV-1 and SARS-CoV-2 RBD structure. **A and B show the** sequence of the SARS-CoV-1 and SARS-CoV-2 RBD with all cysteines highlighted in yellow. The disulfide bridges are indicated as lines between the cysteines in the RBD sequences. These bridges shield parts of the folded protein structure from force and thereby restricts unfolding. The parts of the protein still able to be under force are highlighted in green, shielded parts in grey. The sections under force add up to 51 amino acids (aa, 19 nm for 0.365 nm/aa) for the RBD of SARS-CoV-1 and 54 aa (20 nm for 0.365 nm/aa) for the RBD of SARS-CoV-2 (these include all folded residues as captured in the crystal structure). Some parts of the N-terminus of the RBD used in the tethered ligand protein are probably not folded but will also get released together with the linker increment. The unfolded parts on the C-terminal probably get stretched already in the initial stretching of the linkers for attachment and therefore will not contribute to the length released during the (partial) unfolding of the RBD. In **C** and **D** the corresponding RBDs of SARS-CoV-1 (PDB ID: 2dd8) and SARS-CoV-2 (PDB ID: 6m0j) are depicted, using the same color code as in A,B for parts of the protein under force (green), contributing to the increments observed, and parts shielded from unfolding (grey). The orange circles mark the N- and C-terminal end of the crystal structure whereas the sequence in (A,B) show the entire RBD sequence used in the tethered ligand protein.

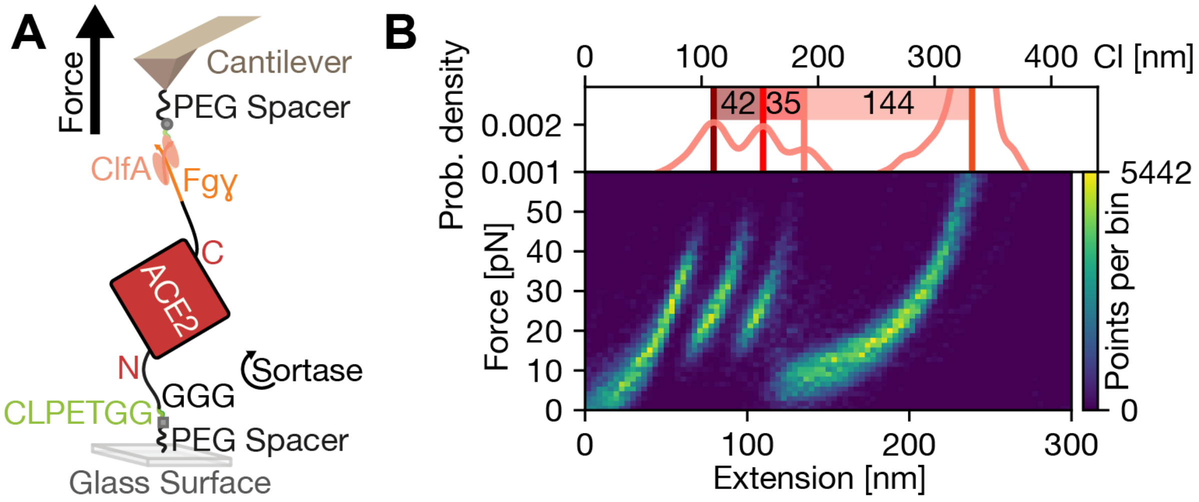


**Supplementary Fig. 3** Unfolding of ACE2 by AFM SMFS. **A** Schematic of the experimental setup pulling on ACE2 which is the same as for the tethered ligand proteins. ACE2 is coupled covalently on a aminosilanized glass surface using heterobifunctional polyethylene glycol (PEG) spacers. The construct is attached in a sortase mediated reaction to a CLPETGG peptide attached to the maleimide of the PEG spacer. For reversible tethering the Fgɣ tag on the tethered ligand protein can be pulled by a ClfA handle. ClfA is covalently attached to the cantilever by an sfp mediated reaction, connecting the ybbR tag to a CoA coupled to a PEG spacer on the AFM cantilever. **B** Heatmap of AFM unfolding traces of ACE2. The heatmap of an overlay of 152 aligned curves shows the characteristic ACE2 trident shaped pattern also observed in the full unfolding of the tethered ligand protein. On top an alignment of all contour length transformed density curves is shown. The contour length increments of the ectodomain ACE2 match well with the last increments of the full tethered ligand protein. This allows the assignment of the last three peaks before the final rupture in the complete RBD:ACE2 tethered ligand protein to the unfolding of the ACE2 ectodomain. This unfolding pattern can be used as a fingerprint for identifying single-molecule traces and normalizing force distributions in measurements if they were recorded with different cantilevers.

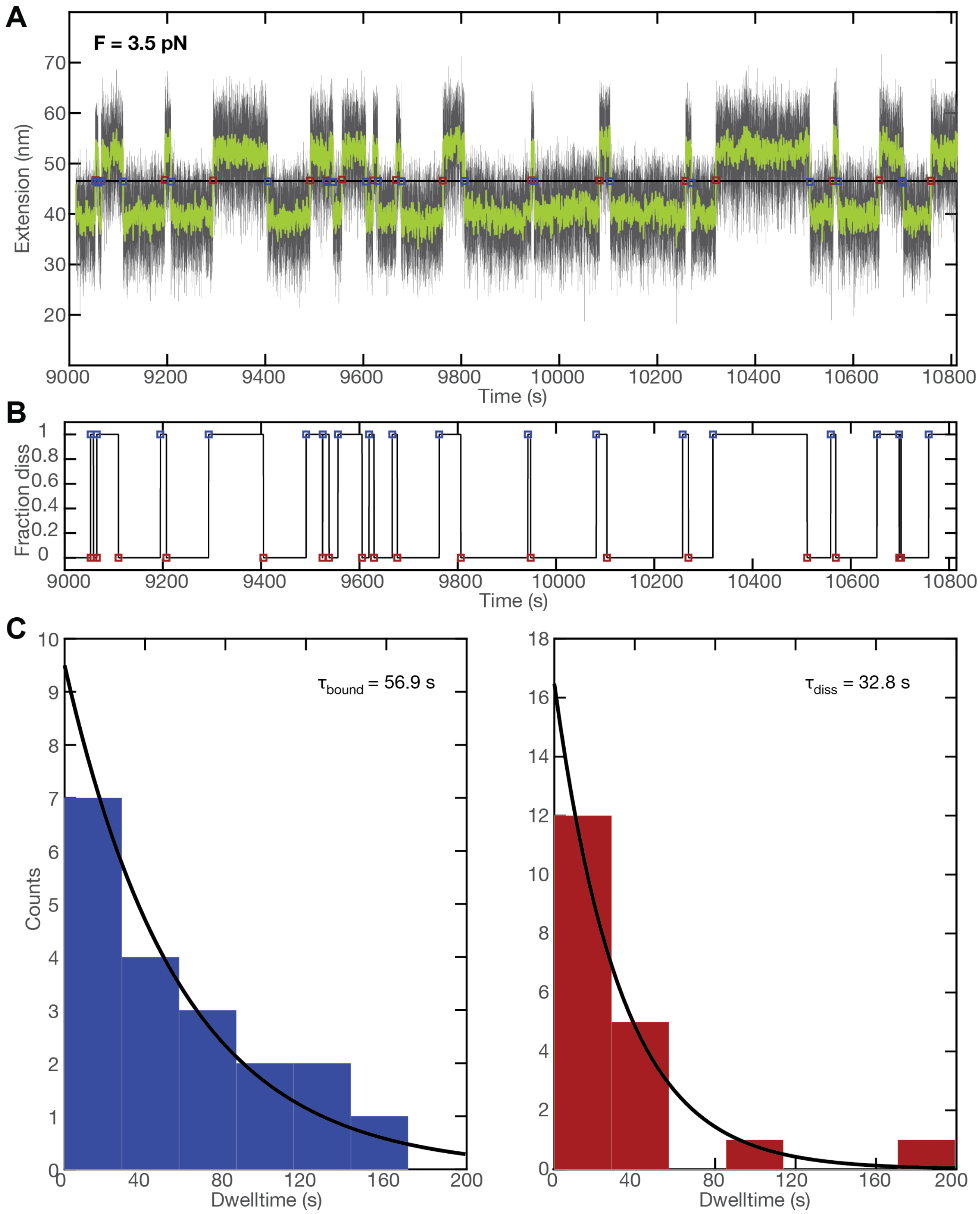


**Supplementary Fig. 4.** Dwell time analysis of the tethered ligand extension time traces in MT. **A** Short segment of an extension time trace measured for an SARS-CoV-2 RBD:ACE2 tethered ligand construct at a stretching force of 3.5 pN. Raw data at 72 Hz are shown in black and filtered data (50 frame moving average) are shown in green. Assignment of the dwell times is based on the filtered data. The black horizontal line is the threshold; blue squares indicate the first data point after crossing the threshold from below, i.e. transition from the bound to the dissociated state; red squares indicate the first data point after crossing the threshold from above, i.e. transition from the dissociated to the bound state. **B** Time trace derived from the analysis shown in panel A, indicating the current state of the tethered-ligand system with “1” corresponding to the dissociated state and “0” to the bound state. The time between the transitions between “0” and “1” correspond to the dwell times. **C, D** Histograms of dwell times in the bound state (**C**) and dissociated state (**D**) obtained from the analysis shown in panels **A** and **B**. The dwell times are well described by single exponential fits, shown as solid lines. Insets show the mean dwell times from maximum likelihood fits of the single exponentials.

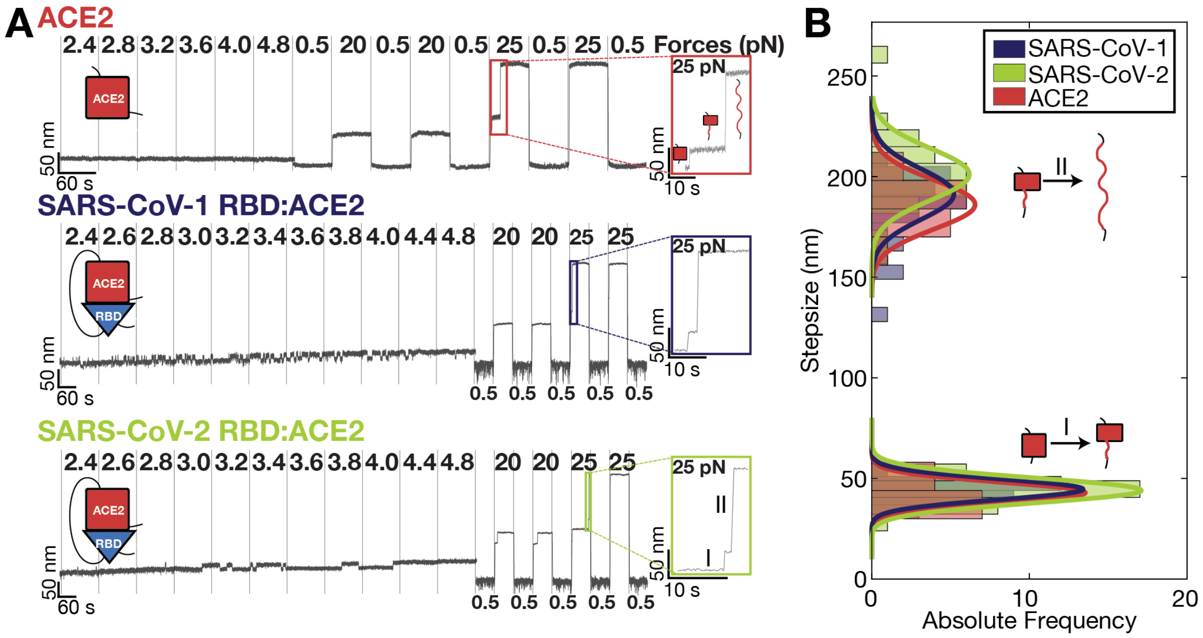


**Supplementary Fig. 5.** ACE2 unfolding events observed in different constructs in the MT. **A** Short force ramps and unfolding jumps for single ACE2 (top), SARS-CoV-1 RBD:ACE2 (middle), and SARS-CoV-2 RBD:ACE2 (bottom), measured in MT. Both tethered ligand constructs show equilibrium hopping transitions in the range between 2.4 and 4.8 pN, corresponding to interface opening and partial RBD unfolding. It is apparent that the transition are much more rapid, i.e. exhibit shorter dwell times, for SARS-CoV-1 compared to SAXS-CoV-2. During the force jumps, all constructs show characteristic two-step unfolding, marking unfolding of the ACE2 domains. **B** Histogram of jump-size distribution of the 2 step-unfolding events, after contour length transformation (with Lp = 0.5 nm) for all constructs shown in **A**. Color codes correspond to color codes in **A**. Solid lines represent gauss fits to the histograms. Distributions agree very well among the different constructs.
